# Quantification of subcellular RNA localization through direct detection of RNA oxidation

**DOI:** 10.1101/2024.11.12.623278

**Authors:** Hei-Yong G. Lo, Raeann Goering, Agnese Kocere, Joelle Lo, Megan C. Pockalny, Laura K. White, Haydee Ramirez, Abraham Martinez, Seth Jacobson, Robert C. Spitale, Chad G. Pearson, Marino J. E. Resendiz, Christian Mosimann, J. Matthew Taliaferro

## Abstract

Across cell types and organisms, thousands of RNAs display asymmetric subcellular distributions. The study of this process often requires quantifying abundances of specific RNAs at precise subcellular locations. To analyze subcellular transcriptomes, multiple proximity-based techniques have been developed in which RNAs near a localized bait protein are specifically labeled, facilitating their biotinylation and purification. However, these complex methods are often laborious and require expensive enrichment reagents. To streamline the analysis of localized RNA populations, we developed Oxidation-Induced Nucleotide Conversion sequencing (OINC-seq). In OINC-seq, RNAs near a genetically encoded, localized bait protein are specifically oxidized in a photo-controllable manner. These oxidation events are then directly detected and quantified using high-throughput sequencing and our software package, PIGPEN, without the need for biotin-mediated enrichment. We demonstrate that OINC-seq can induce and quantify RNA oxidation with high specificity in a dose- and light-dependent manner. We further show the spatial specificity of OINC-seq by using it to quantify subcellular transcriptomes associated with the cytoplasm, ER, nucleus, and the inner and outer membranes of mitochondria. Finally, using transgenic zebrafish, we demonstrate that OINC-seq allows proximity-mediated RNA labeling in live animals. In sum, OINC-seq together with PIGPEN provide an accessible workflow for the analysis of localized RNAs across different biological systems.

## INTRODUCTION

The localization of specific RNA molecules to distinct subcellular locations is a widespread phenomenon. In organisms ranging from yeast to human, thousands of different RNA species are asymmetrically localized in a variety of conditions and cell types (1–4). In many cases, the transport of these RNAs is mediated by the interaction of specific cis-localization-regulatory sequences with RNA-binding proteins (RBPs) that recognize these sequence elements. However, for the vast majority of localized RNAs, the identity of the cis-elements and trans-factors that regulate their subcellular distribution are unknown (5).

Misregulation of subcellular RNA localization is associated with a variety of cellular and organismal phenotypes. Mating type switching in yeast (1) and developmental patterning in Drosophila (6) require proper trafficking of specific RNAs. In humans, a variety of human neurological diseases are linked to defects in neuronal RNA localization (7).

A key challenge in studying RNA localization is identifying which transcripts are enriched at a given subcellular location. Methods to interrogate this problem have been consistently improving over the last few decades. Historically, microscopy-based methods like single molecule fluorescence in situ hybridization (smFISH) have provided high-resolution quantification of localized abundances of single transcripts. These methods have more recently been multiplexed to allow the interrogation of multiple transcripts at once (8, 9). However, microscopy-based techniques require highly sophisticated imaging setups, and the number of cells that can be assayed in a single experiment remains limited. Given that gene expression at the single cell level can be stochastic and prone to “bursts” (10), gathering population average measurements from millions of cells can sometimes be preferable.

Other methods use high-throughput sequencing for the quantification of subcellular transcriptomes. Cells with extended morphologies, including neurons, can be mechanically fractionated into distinct subcellular fractions representing cell bodies and projections (11). RNA collected from these fractions can be analyzed using high-throughput RNA sequencing to identify transcripts that are differentially abundant between the two subcellular locations (12–17). This approach is limited to cells with specific morphologies with elongated projections. Given that RNAs are also localized in cells without these structural features (3, 18), more flexible techniques are needed.

Biochemical subcellular fractionation techniques involving ultracentrifugation have been used (19, 20), yet these approaches separate molecules based on biochemical properties, not necessarily subcellular location. Two molecules that are not spatially coincident in cells that have similar biochemical properties may cofractionate, leading to false positives. Similarly, two molecules that are spatially coincident may not copurify, leading to false negatives.

To improve on these approaches, proximity-based approaches for identifying localized RNAs have been developed (21–24). These approaches rely on a localized “bait” protein and use spatially restricted chemical reactions enabled by the bait protein to specifically label nearby RNAs. Most commonly, labeling events facilitate biotinylation of the bait-proximal RNA, allowing its purification with streptavidin-coated beads and analysis using high-throughput sequencing. These approaches are more flexible in that they can target many subcellular locations in a variety of cultured cell types. However, proximity labeling-based approaches often take multiple days to complete, require relatively expensive RNA enrichment reagents, and due to harsh chemical treatments are often not compatible with experimentation in live organisms.

To streamline the process of subcellular RNA quantification, we developed OINC-seq. Our approach builds on the principles of a previously established proximity-based technique, Halo-seq (23, 25). In Halo-seq, a HaloTag protein domain (26) is genetically fused to a protein that is specifically localized to the subcellular location of interest. A Halo ligand, Halo-dibromofluorescein (Halo-DBF), is then added, which specifically and covalently binds the HaloTag domain, thus assuming the same subcellular distribution as the localized protein. Upon irradiation with green light, Halo-DBF emits singlet oxygen radicals that oxidize biomolecules within a radius of approximately 100 nm. Oxidized (i.e. localized) RNA is then a substrate for in situ alkynylation using the small nucleophile propargylamine. This renders the localized RNA a substrate for in vitro biotinylation using “Click” chemistry (27) and eventual purification using streptavidin.

With OINC-seq, the laborious and expensive biotinylation and purification steps are removed. Instead, in situ RNA oxidation events are read out directly using high-throughput sequencing. Oxidation events are detected as misincorporation at guanosine residues by reverse transcriptase. These oxidation events are then quantified at the individual transcript and gene levels using the accompanying Pipeline for the Identification of Guanosine Positions Erroneously Notated (PIGPEN) software. We demonstrate that OINC-seq induces and quantifies RNA oxidation events in a dose- and light-dependent manner. We show that OINC-seq performs these tasks in a spatially restricted manner, allowing the quantification of RNA populations at specific subcellular locations. Finally, we demonstrate that OINC-seq is compatible with RNA labeling in live zebrafish embryos using transgenic HaloTag fusions. Taken together, as a genetically controllable system with in vivo functionality, OINC-Seq expands the reach of proximity-based RNA localization techniques.

## RESULTS

### 8-oxoguanosine is interpreted by Superscript IV reverse transcriptase as uridine approximately 80% of the time

Of the four nucleosides, guanosine has the lowest reduction potential (28), making it the most likely nucleoside to be oxidized. Oxidation converts guanosine to 8-oxoguanosine (8OG). Further oxidation of 8OG leads to the production of 5-guanidinohydantoin (GH) and spiroiminodihydantoin (**Figure 1A**). Using primer extension and gel electrophoresis, previous work has demonstrated that when reverse transcriptase (RT) encounters these lesions, it makes predictable incorporation errors at those locations in the resulting cDNA (29). Specifically, RT had been observed to interpret 8OG as uridine approximately 60% of the time while further oxidized products were interpreted as either cytidine or uridine (29). After PCR of the resulting cDNA, these misincorporations are manifested as G to T and G to C mutations in the strand corresponding to the original RNA molecule.

**Figure 1:**
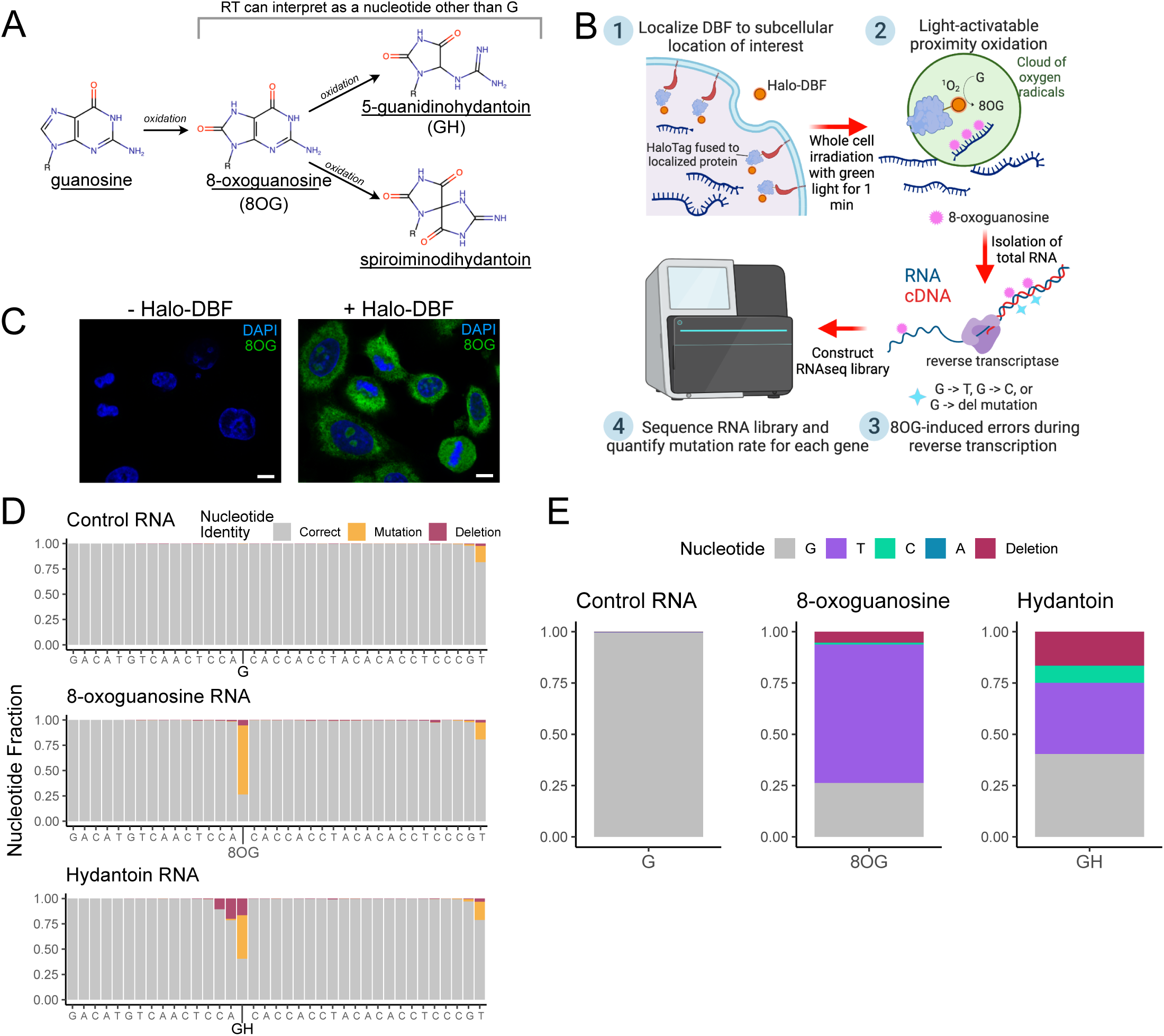
OINC-seq detects oxidized RNA by quantifying reverse transcriptase errors induced by guanosine oxidation. (A) Schematic showing the oxidative products of guanosine. (B) Schematic of an OINC-seq experiment. Guanosine oxidation, induced by proximity-mediated labeling, results in RT misincorporation events. These events are read out using high-throughput sequencing as G to T mutations, G to C mutations, and G deletions relative to a reference sequence. (C) Immunofluorescence detection of 8OG in HeLa cells expressing a cytoplasmic p65 Halo fusion protein with (+Halo-DBF) or without labeling (-Halo-DBF). Scale bars 8 µm. (D) Frequency of nucleotide conversions (orange) and deletions (red) in an RT-PCR product of a synthesized RNA oligonucleotide containing a control guanosine residue at position 16 (top), a single 8OG residue at position 16 (middle) or a single GH residue at position 16 (bottom). (E) Proportion of nucleotide content of RT-PCR products at position 16 across all reads for the control oligonucleotide, 8OG-containing oligonucleotide, or GH-containing oligonucleotide.

In Halo-seq, bait-proximal RNAs are oxidized and alkynylated *in situ*, facilitating their biotinylation *in vitro* and eventual enrichment with streptavidin (25). We wondered whether we could instead harness RT’s propensity to misinterpret oxidized guanosine residues to directly quantify the amount of a given RNA species that had been HaloTag-proximal in cells, potentially negating the need for biotinylation and enrichment. We reasoned that this could be done by isolating whole-cell RNA after the light-controlled oxidation event, identifying the resulting modified RNA species using RNAseq, and quantifying RNA oxidation events by counting G to T and G to C conversions on a per-gene basis (**Figure 1B**).

For this strategy to work, oxidative labeling produced using Halo-DBF must produce 8OG residues in cells. To test this, we ectopically expressed a cytoplasmically localized Halo-tagged protein (Halo-p65) in HeLa cells. We then added the light-sensitive Halo ligand Halo-DBF (23), and irradiated the cells with green light for 5 minutes to induce RNA oxidation (see **Methods** for details). Using an antibody that targets 8OG, we measured the abundance of 8OG in this sample and a control sample in which we omitted the addition of Halo-DBF. We observed an 18-fold increase in 8OG abundance in Halo-DBF-treated cells, indicating that radicals produced through optical activation of localized Halo-DBF were creating 8OG residues (**Figure 1C, S1A**).

We next moved to the detection of 8OG using sequencing-based approaches. Although RT had been previously shown to misinterpret 8OG approximately 60% of the time (29), this rate was calculated using primer extension and gel electrophoresis assays that are challenging to accurately quantify. We therefore sought to establish a more quantitative understanding of the frequency at which 8OG-caused misinterpretation of guanosine by RT occurs. To do this, we synthesized two RNA oligonucleotides of identical sequences with one exception. In one oligonucleotide, a single 8OG residue was inserted at a known nucleotide position. At the corresponding position in the control oligonucleotide, there was a guanosine (**Figure S1B-E**). We then reverse transcribed these oligos using Superscript IV, amplified the resulting cDNA, and sequenced it using an Illumina-based platform.

In our sequencing results, we found that at most positions in both oligos, nucleotide identities in the amplified cDNA were as expected almost 100% of the time. However, in the 8OG-containing oligo, the 8OG position was replaced with a nucleotide other than guanosine approximately 70% of the time (**Figure 1D**). Further inspection of this position showed that almost all of these mutations were G to T mutations, consistent with the previously reported ability of Superscript-based RTs to interpret 8OG as uridine (**Figure 1E**) (29). We repeated this experiment with an RNA oligo containing a further oxidized guanosine product (5-guanidinohydantoin or GH). We observed conversions at the nucleotide substituted with GH 50% of the time (**Figure 1D**). We found that almost all of these conversions were G to T and G to C conversions (**Figure 1E**, **S1F**), as previously shown (29). In our data, G to T conversions are therefore likely arising from both 8OG and hydantoin guanosine oxidation products while G to C conversions are arising only from hydantoins. Interestingly, we also observed that RT also had a propensity to skip oxidized guanosine residues, particularly the hydantoin, resulting in a deletion of the nucleotide in cDNA (**Figure 1E**). This effect had not been previously reported.

These results gave us confidence that Superscript IV would be suitable for 8OG quantification. Since this technique leverages nucleotide conversions (mutations) for the study of RNA oxidation, we named it Oxidation-Induced Nucleotide Conversion Sequencing (OINC-Seq).

### PIGPEN quantifies mutations in RNAseq data

Given that we could quantify 8OG residues in single RNA oligos (**Figure 1D, E**), we then moved to testing 8OG residues created through localized oxidation in cells. To facilitate this, we created software that can identify nucleotide conversions from RNAseq data. With this approach, we can identify and quantify conversions induced by guanosine oxidation (i.e. G to T, G to C, and G deletion events, hereafter referred to as G conversions) in RNAseq samples. We named this software the Pipeline for the Identification of Guanosine Positions Erroneously Notated (PIGPEN) (**Figure 2A**).

**Figure 2:**
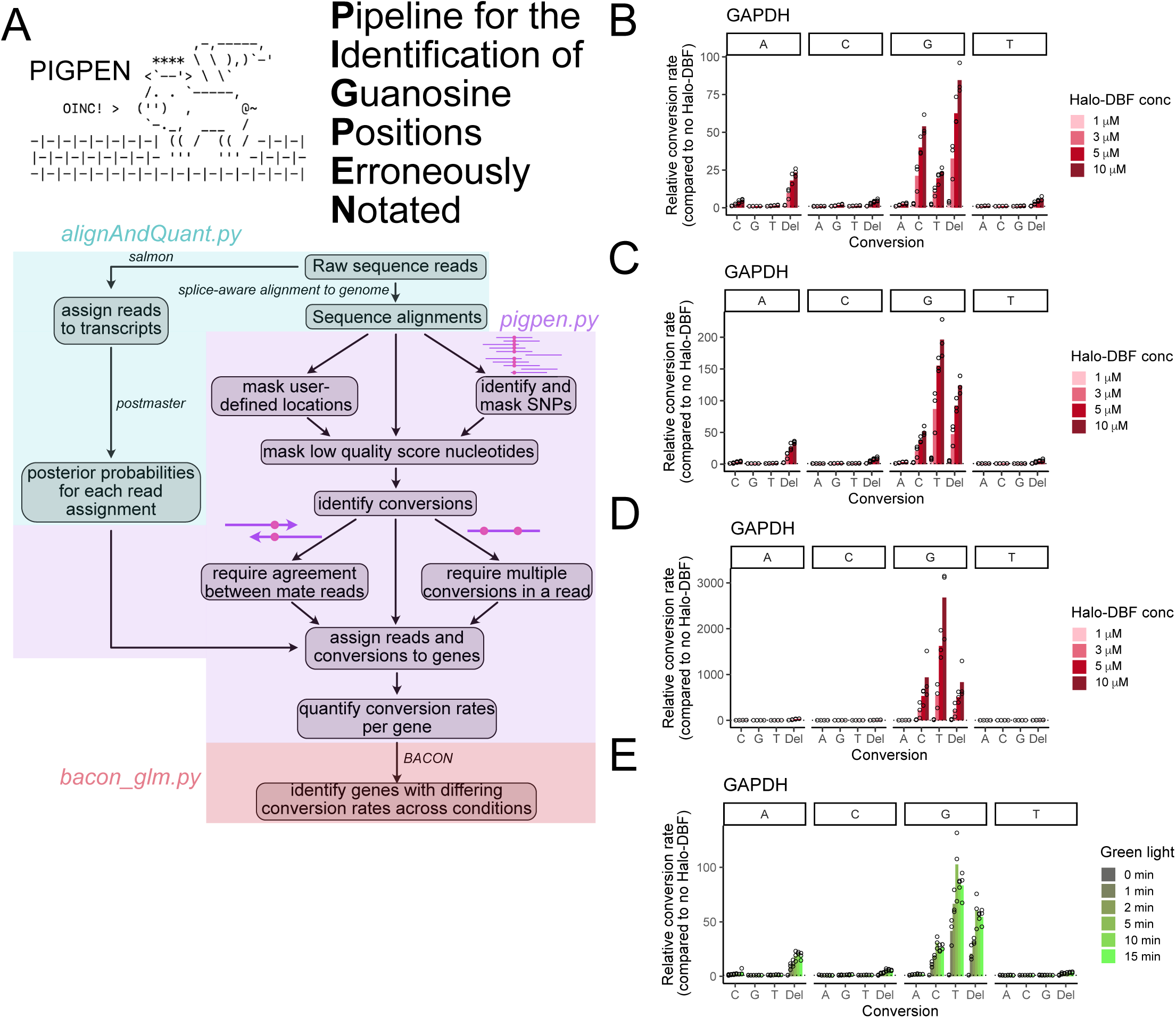
PIGPEN detects nucleotide conversions in RNAseq data and quantifies proximity-induced RNA oxidation events in OINC-seq experiments. (A) Schematic of PIGPEN workflow. (B-D) Proximity-mediated RNA oxidation was induced using a cytoplasmically-localized HaloTag protein and increasing amounts of the oxygen radical-producing Halo-DBF ligand. Relative conversion rates in an RT-PCR amplicon of the GAPDH transcript were calculated and compared to samples in which Halo-DBF was omitted (n=3). Multiple analyses were performed with varying PIGPEN parameters, including whether one G conversion was required to be found in a single read of a mate pair (B), one conversion was required in both reads of a mate pair (C), or two conversions were required per read (D). (E) Proximity-mediated RNA oxidation was induced using a cytoplasmically-localized HaloTag protein and 5μM Halo-DBF. Samples were then irradiated with green light for increasing amounts of time. Nucleotide conversion rates were then calculated using PIGPEN and compared to samples in which Halo-DBF was omitted. In this analysis, only one G conversion was required, and conversions were required to be found in both mates of a read pair.

In brief, PIGPEN first aligns input RNAseq reads to a genome and then calls mutations relative to the reference sequence. Optionally, reference positions known to contain high frequencies of mutations (i.e. single nucleotide polymorphisms (SNPs)) can be automatically called and masked from further analysis. Further, to minimize the contribution of sequencing errors, a requirement for dual observation of a given mutation in both mates of a read pair can be enforced.

In parallel, reads are assigned to transcripts using the quasi-mapping software salmon (30). By combining observed conversions within reads with their salmon-based assignments, conversions are then (fractionally) assigned to transcripts. Gene-level conversion counts are then obtained by summing transcript-level conversion counts across all transcripts assigned to a given gene. Finally, optionally, genes whose G conversion rates significantly differ across conditions can be identified using the statistical framework in the Bioinformatic Analysis of the Conversion of Nucleotides (BACON) module.

To test the ability of PIGPEN to accurately quantify mutation rates in RNAseq data, we created a simulated paired-end RNAseq dataset in which all twelve possible inter-nucleotide conversion rates were predefined. On top of these conversion rates, we also simulated a sequencing error rate of 0.001, which is the approximate rate of sequencing error on Illumina platforms (31). We found that under these conditions, PIGPEN accurately calculated nucleotide conversion rates that were above the sequencing error rate, but that any conversion rates below the sequencing error rate were effectively dominated by sequencing error (**Figure S2A, left**). However, if we required that observed conversions be present in both mates of a read pair, sequencing error was greatly reduced, and nucleotide conversion rates were accurately quantified across a range of frequencies (**Figure S2A, right**).

To test the ability of PIGPEN to identify and quantify 8OG sites in transcriptome-scale RNA-seq data, we turned to a dataset in which 8OG-containing transcripts had been enriched using the anti-8OG antibody we used previously for immunofluorescence (**Figure 1C**) (32). In this dataset, oxidized RNAs were identified by comparing transcript abundances in input and immunoprecipitated RNA samples. Using PIGPEN, we found a 50% higher rate of G to T conversions in the immunoprecipitated RNA compared to the input, consistent with an increased abundance of 8OG in these samples (**Figure S2B, C**). Conversely, G to C conversions and G deletions were only ∼10% higher in the immunoprecipitated RNA, consistent with the antibody recognizing 8OG and not further oxidized species (**Figure 1A, E**). Together, these simulated and real-life data analyses establish PIGPEN as a software pipeline to quantify 8OG frequencies in RNAseq data.

### OINC-seq conversions are Halo-DBF dose-dependent

Given that we could detect and quantify 8OG residues in RNAseq data, we then moved to quantifying their induced formation from the localized oxidation reactions used by RNA proximity labeling experiments. We utilized our HeLa line that inducibly expressed a broadly cytoplasmically localized Halo-tagged protein, Halo-p65 (**Figure S2D**). We then added increasing amounts of Halo-DBF to these cells, washed away unbound Halo-DBF, irradiated them with green light for 5 minutes, and immediately collected total RNA samples. As a control, we also collected untreated samples without Halo-DBF addition. From all samples, we analyzed an amplicon from the *GAPDH* transcript using RNAseq and used PIGPEN to quantify nucleotide conversions in the resulting reads.

Compared to untreated samples in which Halo-DBF was omitted, we saw a large (up to 100 fold), dose-dependent increase in G conversions in Halo-DBF-treated samples, indicative of the formation and detection of guanosine oxidation products (**Figure 2B**). Of the other 13 possible nucleotide conversions, only A deletions were similarly enriched in a dose-dependent manner, demonstrating the specificity of the reaction.

As mentioned above, PIGPEN optionally only considers conversions observed in both mates of a read pair. We quantified conversion rates with (**Figure 2C, S2E**) and without (**Figure 2B**) this option. Requiring conversions to be present in both mates significantly increased conversion enrichment from 100 fold to 200 fold, suggesting that requiring agreement between both read pairs is effective in reducing the contribution of Illumina-based sequencing errors.

Although G conversions are suggestive of 8OG residues, these conversions also exist in non-Halo-DBF treated samples due to reasons unrelated to Halo-DBF-dependent oxidation (e.g., infidelity of the reverse transcriptase, naturally occurring 8OG, etc.). These “non-specific” conversions may be expected to be randomly scattered across all the guanosine residues in an RNA sample. On the other hand, due to the proximity-dependent nature of the reaction, we expect that Halo-DBF-dependent conversions are more likely to be spatially clustered within the cell and therefore found together within a single RNA molecule. We therefore tested whether we could increase signal to noise levels for proximity-dependent 8OG creation by only counting G conversions that came from RNAseq reads with at least two of these conversions within the ∼300 bp that were interrogated in the amplicon. In fact, this approach further increased conversion enrichment levels considerably, with G to C conversions now up to 3000-fold enriched in Halo-DBF-treated samples compared to untreated samples (**Figure 2D**). The dose-dependent relationship between Halo-DBF concentration and conversion rates remained, again demonstrating that these nucleotide conversions were likely due to proximity-dependent RNA oxidation events.

In HeLa cells, *GAPDH* transcript is very highly expressed at approximately 1000 tpm (transcripts per million). To demonstrate the ability of OINC-seq to quantify RNA oxidation on more moderately expressed transcripts, we targeted an amplicon of the *CD9* gene. Transcripts from this gene are 20-fold more lowly expressed than those from *GAPDH*. We again found that G conversion rates were approximately 100 folder higher in Halo-DBF treated cells compared to untreated cells (**Figure S2F**).

### OINC-seq mutations are dependent upon irradiation with green light

A key feature of a proximity labeling experiment is temporal control as this lends additional specificity to the experiment. In Halo-seq and OINC-seq, this is achieved through photo-activation of the Halo-DBF ligand with green light. To assay the photo-dependence of the OINC-seq reaction, we performed proximity labeling in cells expressing Halo-p65 using increasing exposure times to green light (see **Methods** for details). To quantify guanosine oxidation, we again analyzed an amplicon of the *GAPDH* transcript.

We found that samples that were not exposed to green light did not accumulate G conversions above background (**Figure 2E**). Increasing amounts of green light exposure time led to increasing rates of G conversions, with the accumulation plateauing at 5 minutes (**Figure 2E, S2G**). We therefore concluded that the OINC-seq reaction was dependent upon exposure to green light and that for this Halo fusion and RNA pair, the reaction was saturated at 5 minutes of green light exposure. We believe this rather short period of labeling minimizes oxidative damage and secondary, off-target effects.

### Testing the misincorporation frequency of different reverse transcriptases at 8OG residues

OINC-seq relies on the ability of reverse transcriptase to interpret 8OG residues as a nucleotide other than guanosine. Up to this point, we had used Superscript IV RT for this purpose as we and others had observed that this enzyme had this property (29). We sought to test if other reverse transcriptases may perform misincorporation at 8OG sites at higher rates and therefore be a superior choice for OINC-seq studies.

We therefore repeated the OINC-seq experiment again using Halo-p65 and a *GAPDH* amplicon. We reverse-transcribed the resulting RNA from this experiment using four different RT enzymes: Superscript IV, AccuScript, MarathonRT, and TGIRT III. We were unable to generate sufficient cDNA product for analysis using MarathonRT and TGIRT III. After quantifying nucleotide conversions with PIGPEN, we found that Superscript IV generated G conversions in the Halo-DBF-treated RNA at a higher rate than AccuScript (**Figure S2H**). We therefore continued to use Superscript IV for all future experiments.

### OINC-seq creates and quantifies oxidized guanosines in a spatially specific manner

We found that OINC-seq was able to induce and quantify RNA oxidation, but in order for the technique to be useful for the study of localized RNAs, the oxidation must happen in a spatially restricted and specific manner. To test the spatial specificity of OINC-seq, we created two HeLa lines, one that inducibly expressed a cytoplasmically localized Halo fusion (Halo-p65) and another that inducibly expressed a nuclearly localized Halo fusion (Halo-H2B) (**Figure 3A**). We then induced proximity-dependent RNA oxidation in both lines and used PIGPEN to quantify oxidized guanosines in amplicons of *MALAT1* and *GAPDH*.

**Figure 3:**
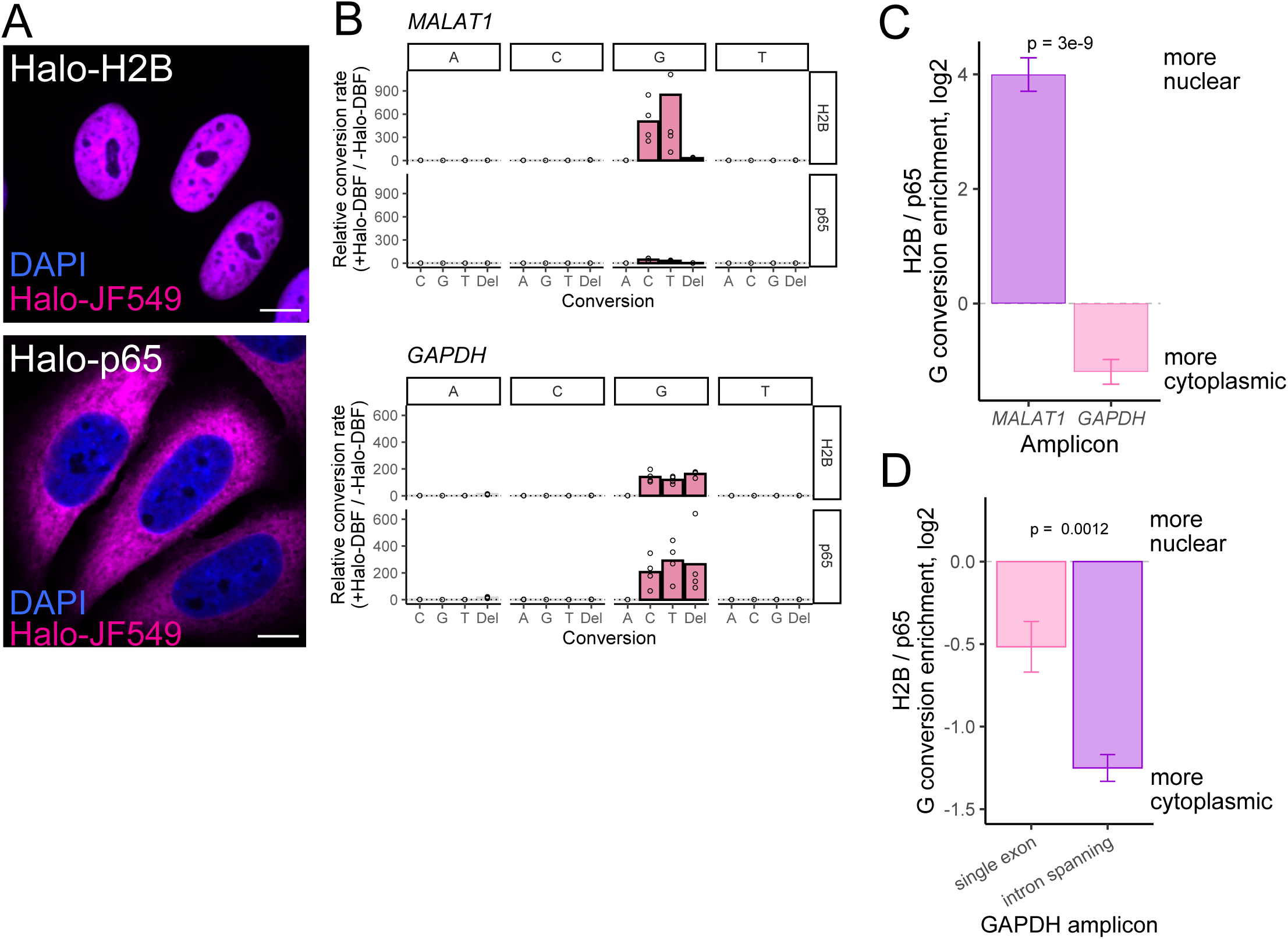
OINC-Seq labels RNA in a spatially specific manner. (A) HaloTag fusions to H2B (top) and p65 (bottom) are nuclearly and cytoplasmically localized, respectively. HaloTag fusions are visualized using a fluorescent Halo ligand, JF549 (magenta). Scale bars 8 µm. (B) Relative nucleotide conversion rates between Halo-DBF treated and untreated samples (n=4). Results are shown for RT-PCR amplicons of MALAT1 (top) and GAPDH (bottom) in cells expressing either Halo-H2B or Halo-p65. In this analysis, conversions were required to be seen in both reads of a mate pair, and reads were required to have at least two G conversions. (C) Relative enrichment of G conversions in Halo-H2B samples compared to Halo-p65 samples. (D) Relative enrichment of G conversions in Halo-H2B samples compared to Halo-p65 samples for a non-intron-spanning (left) and intron-spanning (right) amplicon of GAPDH.

*MALAT1* RNA is strongly enriched in the nucleus (33). *GAPDH* RNA is present in both the nucleus and the cytoplasm, but relative to *MALAT1* is expected to accumulate to a higher level in the cytoplasm at steady state (**Figure S3A**). Therefore, if the OINC-seq labeling were spatially specific, we would expect to see relatively more G conversions in *MALAT1* than *GAPDH* in the nuclear labeling. Conversely, we would expect more conversions in *GAPDH* than *MALAT1* in the cytoplasmic labeling.

For the *MALAT1* amplicon, we found that in the nuclear labeling, G conversions were approximately 500-fold enriched in the Halo-DBF-treated samples compared to untreated samples. However, in the cytoplasmic labeling, these conversions were only approximately 50-fold enriched in Halo-DBF treated samples compared to untreated samples (**Figure 3B, S3B-C**).

For the *GAPDH* amplicon, the situation was reversed. In the nuclear labeling, G conversions were approximately 200-fold enriched, while in the cytoplasmic labeling, they were 300-fold enriched (**Figure 3B, S3D-E**). Nascent *GAPDH* RNA oxidation in the nuclear labeling is expected as the Halo-fusion is chromatin-associated and the amplicon analyzed does not cross an exon-exon boundary.

*MALAT1* RNA was 16-fold more oxidized in the nuclear labeling while *GAPDH* RNA was 1.5-fold more oxidized in the cytoplasmic labeling (**Figure 3C**). Putting these results together, we observed with OINC-seq that *MALAT1* RNA was approximately 24-fold more enriched in the nucleus than *GAPDH* RNA. We conclude that the RNA oxidation induced by OINC-seq and quantified by PIGPEN occurs in a spatially specific manner, making the technique suitable for RNA localization experiments.

To further probe the spatial selectivity of OINC-seq, we again labeled RNA using the H2B and p65 Halo-fusions. We then amplified two different amplicons of *GAPDH* RNA. One amplicon was the same as used above (**Figure 2B-E, 3B-C**) and was entirely contained within one exon. As such, it is expected to be found on both nascent pre-mRNA and spliced mature mRNA. The other amplicon crossed a spliced junction and is therefore only present in spliced mature mRNA. Given that most cytoplasmic transcripts are spliced while those in the nucleus are a mixture of spliced and unspliced, we would expect the spliced amplicon to be relatively less abundant in nuclear RNA. This prediction was borne out by OINC-seq, further demonstrating the ability of the method to label RNA in a spatially specific manner (**Figure 3D**).

### OINC-seq quantifies RNA localization on a transcriptomic scale

Beyond quantification of RNA localization of single transcripts through amplicon analysis, we next sought to apply OINC-seq for transcriptome-wide analysis of RNA localization. To test the ability of OINC-seq to report on the full RNA content of different subcellular locations, we took advantage of four HeLa lines that inducibly expressed HaloTag fusion proteins targeted to different locations. Halo-p65 was broadly cytoplasmically localized, Halo-P450 was localized to the cytoplasmic face of the endoplasmic reticulum (ER), Halo-ATP5MC1 was localized to the mitochondrial matrix, and Halo-H2B was localized to the nucleus (**Figure 4A**).

**Figure 4:**
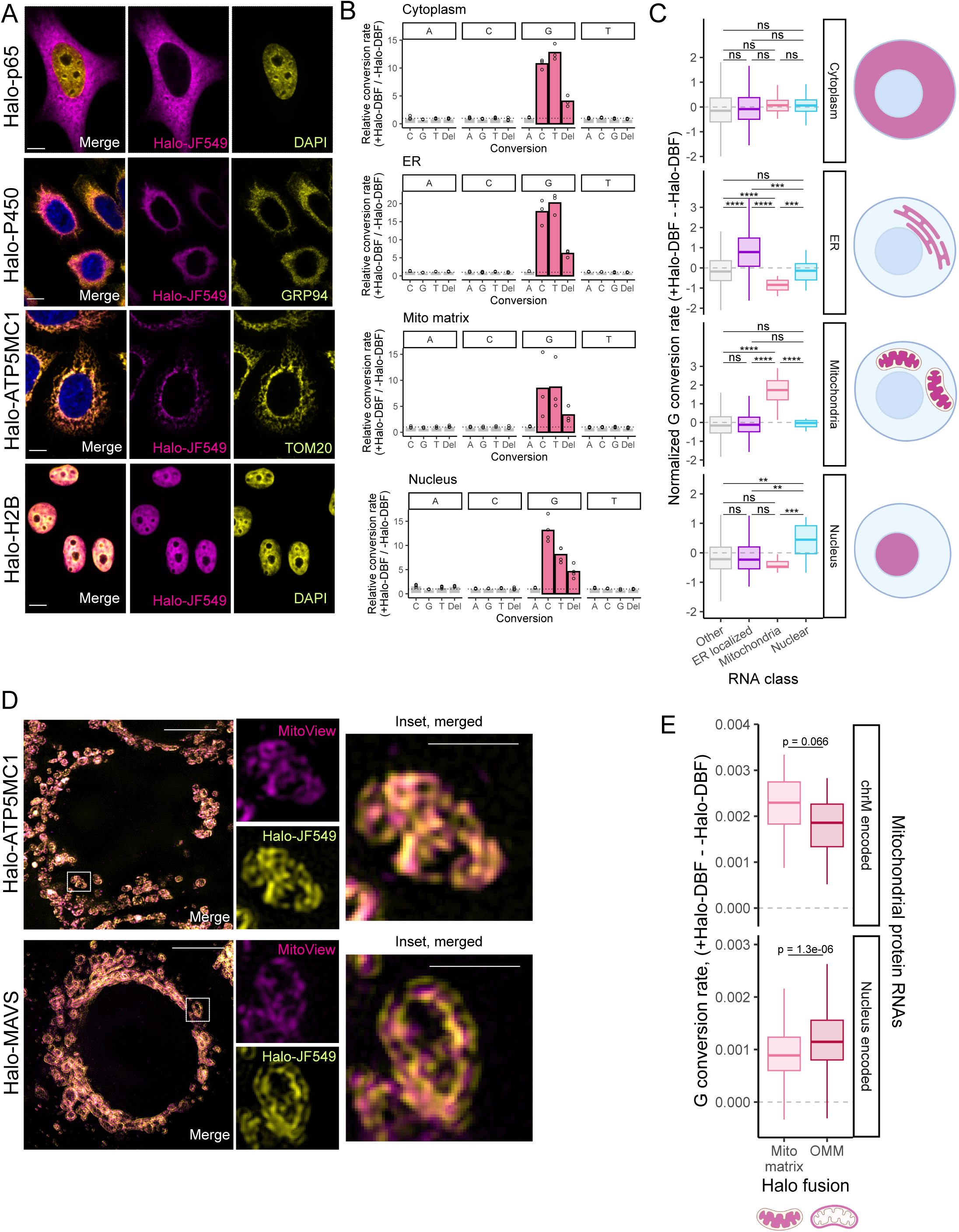
OINC-seq identifies subcellular localized RNAs on a transcriptomic scale. (A) Representative fluorescent images of HaloTag fusion proteins co-localized with proteins at the region of interest. HaloTag fusion proteins are visualized by the Halo ligand JF549 (magenta). Top: Halo-p65. Top-Middle: ER-localized HaloTag fusion Halo-P450 colocalizing with a known ER marker GRP94 (yellow). Bottom-Middle: mitochondrial matrix HaloTag fusion Halo-ATP5MC1 colocalizing with the mitochondrial protein TOM20 (yellow). Bottom: H2B HaloTag fusion Halo-H2B colocalizing with DAPI (yellow). Scale bar 8 µm. (B) Bulk relative conversion rates comparing Halo-DBF treated and untreated samples (n=3) for all four labeled subcellular compartments. (C) G conversion rates of four different classes of RNAs (ER-localized, Mitochondria, Nuclear, Other) in OINC-seq experiments using cytoplasmically-, ER-, mitochondrially-, and nuclearly-localized HaloTag fusions. Wilcox rank-sum tests were used to compare across RNA classes (ns represents p > 0.05, ** p < 0.01, **** p < 0.0001) (D) Representative fluorescent images of mitochondrial-localized HaloTag fusions Halo-ATP5MC1 and Halo-MAVS using structured illumination microscopy. Top: the mitochondrial matrix Halo fusion (Halo-ATP5MC1) co-stained with the mitochondrial membrane marker MitoView. Bottom: the outer mitochondrial membrane Halo fusion (Halo-MAVS) co-stained with the mitochondrial membrane marker MitoView. Scale bar 5 µm. Insets allow better appreciation of the relative position of each stained structure. Scale bar 1 µm (insets). (E) G conversion rates for two classes of RNA that encode mitochondrial proteins, those encoded on the mitochondrial chromosome and those encoded in the nuclear genome. Rates are shown for OINC-seq experiments using the mitochondrial matrix HaloTag fusion (Halo-ATP5MC1) and the outer mitochondrial membrane HaloTag fusion (Halo-MAVS) (n=3). Wilcox rank-sum tests were used to compare across RNA classes.

We performed OINC-seq with each cell line, isolated RNA, and created Quantseq 3′ end RNAseq libraries (including using this kit’s reverse transcriptase) (34). We then quantified RNA oxidation in each sample using PIGPEN. For each HaloTag fusion, G conversions were enriched in Halo-DBF-treated samples compared to untreated samples, indicating that OINC-seq was inducing RNA oxidation events (**Figure 4B**). However, these conversion enrichment values were noticeably lower than those previously observed in *GAPDH* and *MALAT1* amplicons (**Figure 2C**).

We speculated that this could be due to decreased read depth per gene (thousands of reads) in a transcriptome-scale experiment compared to the per-gene depth in a single amplicon experiment (millions of reads). To test this, we downsampled read depths in the *GAPDH* amplicon analysis from Figure 2B to 10,000 reads. This downsampling had essentially no effect on detected conversion rates (**Figure S4A**), indicating that read depth was not responsible for the decreased conversion enrichment values.

We next wondered whether differences in sequencing library preparation techniques (i.e. RT-PCR vs. 3′ end RNAseq libraries) could affect guanosine conversion frequencies. Indeed, when comparing Halo-DBF-untreated samples, Quantseq 3′ end RNAseq libraries showed significantly higher rates of “background” guanosine conversions than was observed with RT-PCR amplicons (**Figure S4B**). We therefore speculate that differences in the fidelity of polymerases used in the RT-PCR and RNAseq library approaches may be behind the drop in G conversion signal in transcriptome-wide samples. Nevertheless, clear ∼10-fold enrichments of G conversions could be seen in Halo-DBF treated samples, so we continued with analysis of these samples.

To assay OINC-seq’s ability to label subcellular RNA populations, we then focused on G conversion rates in RNAs known to be localized to the cytoplasmic surface of the ER and to the mitochondrial matrix. mRNAs that encode secreted or membrane-embedded proteins are often translated on the surface of the ER, and their presence there has been previously assayed (35). We therefore used these mRNAs as positive controls for ER-localized transcripts. Transcripts that arise from the mitochondrial genome reside in the mitochondrial matrix, and we used these RNAs as positive controls for mitochondrially localized transcripts. Many lncRNAs are known to be enriched in the nucleus, and we used these as positive controls for nuclearly localized transcripts.

G conversion rates of individual samples clustered by the subcellular location of the sample’s Halo fusion, indicating reproducible patterns of guanosine oxidation within a cell line and distinct patterns of guanosine oxidation across cell lines (**Figure S4C**). When RNA was labeled using the broadly cytoplasmically localized Halo-p65 fusion, we observed broad labeling of many classes of RNAs (**Figure 4C**). Conversely, when RNA was labeled with the ER-localized Halo-P450 fusion, we found that known ER-localized RNAs had higher rates of G conversions, consistent with OINC-seq being able to specifically label this RNA population (**Figure 4C**). When RNA was labeled with the mitochondrially localized Halo-ATP5MC1 fusion, mitochondrial RNAs were strongly enriched for G conversions (**Figure 4C**). Finally, when RNA was labeled with the nuclearly localized

Halo-H2B sample, known nuclear RNAs were strongly enriched for G conversions. Using BACON, we identified RNAs in the ER, mitochondrial, and nuclear HaloTag samples that displayed statistically significantly different rates of G conversions, again finding that each compartment was significantly associated with positive control RNAs known to reside there (**Figure S4D-F, Supplemental Tables 2-4**). Taken together, these results indicate that OINC-seq has the ability to assay RNA localization on a transcriptomic scale across different subcellular compartments.

### OINC-seq quantifies RNA localization with high spatial specificity across mitochondrial membranes

To more thoroughly test the spatial discrimination possible with OINC-seq, we compared RNA localization patterns across the mitochondrial double membrane. As a mitochondrial matrix marker, we used our Halo-ATP5MC1 fusion (**Figure 4A**). We compared OINC-seq results from this fusion to those from a Halo-MAVS fusion protein that localized to the outer mitochondrial membrane (36) with the HaloTag domain facing towards the cytoplasm (**Figure S4G**). These two HaloTag fusions are therefore in close proximity of each other within cells, with Halo-MAVS on the outer mitochondrial membrane facing outward and Halo-ATP5MC1 on the inner mitochondrial membrane facing inward.

We verified the localization of these fusion proteins using fluorescent Halo ligands, a mitochondrial dye (MitoView Green), and Structured Illumination (SIM) microscopy. We found that Halo-ATP5MC1 signal consistently coincided with mitochondrial signal (**Figure 4D, top**), consistent with this fusion being localized to the mitochondrial matrix. Conversely, Halo-MAVS signal surrounded the mitochondrial signal, consistent with its localization to the outer mitochondrial membrane (**Figure 4D, bottom; Figure S4G-H**).

Mitochondrial protein-encoding transcripts from the nuclear genome are often translated on the outer surface of mitochondria to facilitate co-translational import of the nascent protein (37). If OINC-seq were able to spatially discriminate RNA localization across the mitochondrial membranes, then we would expect these nuclear-encoded mitochondrial RNAs to be relatively more labeled with the Halo-MAVS fusion while transcripts that arise from the mitochondrial genome should be relatively more labeled with the Halo-ATP5MC1 fusion.

Using amplicons of two mitochondrially-encoded genes, we observed this to be the case (**Figure S4I**). We then expanded this analysis transcriptome-wide. Nucleus-encoded mitochondrial RNAs were indeed more labeled with the Halo-MAVS outer membrane fusion. On the other hand, those encoded by the mitochondrial genome were more labeled with the Halo-ATP5MC1 mitochondrial matrix fusion, although not statistically significantly so (**Figure 4E**). Taken together, genetically encoded HaloTags enable highly sensitive spatial resolution of RNAs using OINC-seq.

### OINC-seq labels RNA in live animals

To date and to our knowledge, all reported RNA proximity labeling techniques have been performed in cultured cells (21–24, 38). Applying RNA proximity labeling into animal models *in vivo* is challenging due to the toxicity of some reagents, particularly those involved in biotinylation, and their inability to penetrate deep into tissues. In contrast, OINC-seq circumvents the need for biotinylation. We therefore sought to perform proximity-based oxidation of RNA in live zebrafish embryos.

For this purpose, we created the Tol2-based transgenic zebrafish strain *Tg(bact2:HaloTag-NES-2A-mCerulean-CAAX)*, or *bact2:HaloTag-NES* for short. The transgene consists of the broadly active, strong *beta-actin2* (*bact2*) promoter driving a HaloTag domain fused to three copies of the nuclear export sequence (NES) from MAPPK2 followed by P2A-membrane-bound mCerulean to identify transgenic zebrafish by blue fluorescence (**Figure S5A, top**). Additionally, we made a similar transgenic line expressing a Halo-H2B fusion (*bact2:HaloTag-H2B*). (**Figure S5A, bottom**). Through standard Tol2-based transgenesis, we isolated and selected a transgenic insertion that showed 50% Mendelian segregation indicative of single-copy transgenes as detectable by ubiquitous mCerulean fluorescence throughout the first 5 days of development (**Figure S5B-C**).

To verify functionality of the HaloTag fusion expressed by the *bact2:HaloTag-NES* transgene, we incubated 2 days post fertilization (dpf) zebrafish embryos homozygous for the transgene with a far-red fluorescent Halo ligand, Halo-JF646. We then washed out unbound ligand and imaged the incubated embryos using confocal fluorescence microscopy (**Figure S5D-E**). In Halo-JF646-treated wild type embryos, we observed only background fluorescent signal. Conversely, in Halo-JF646-treated transgenic zebrafish, we detected fluorescent signal throughout the embryo, indicating expression of the HaloTag-containing transgene (**Figure 5A, S5D-E**). Higher magnification revealed fluorescent signal concentrated in the cytoplasm and nucleus for the *bact2:HaloTag-NES and bact2:HaloTag-H2B* lines, respectively, consistent with their predicted subcellular localizations (**Figure 5B**).

**Figure 5:**
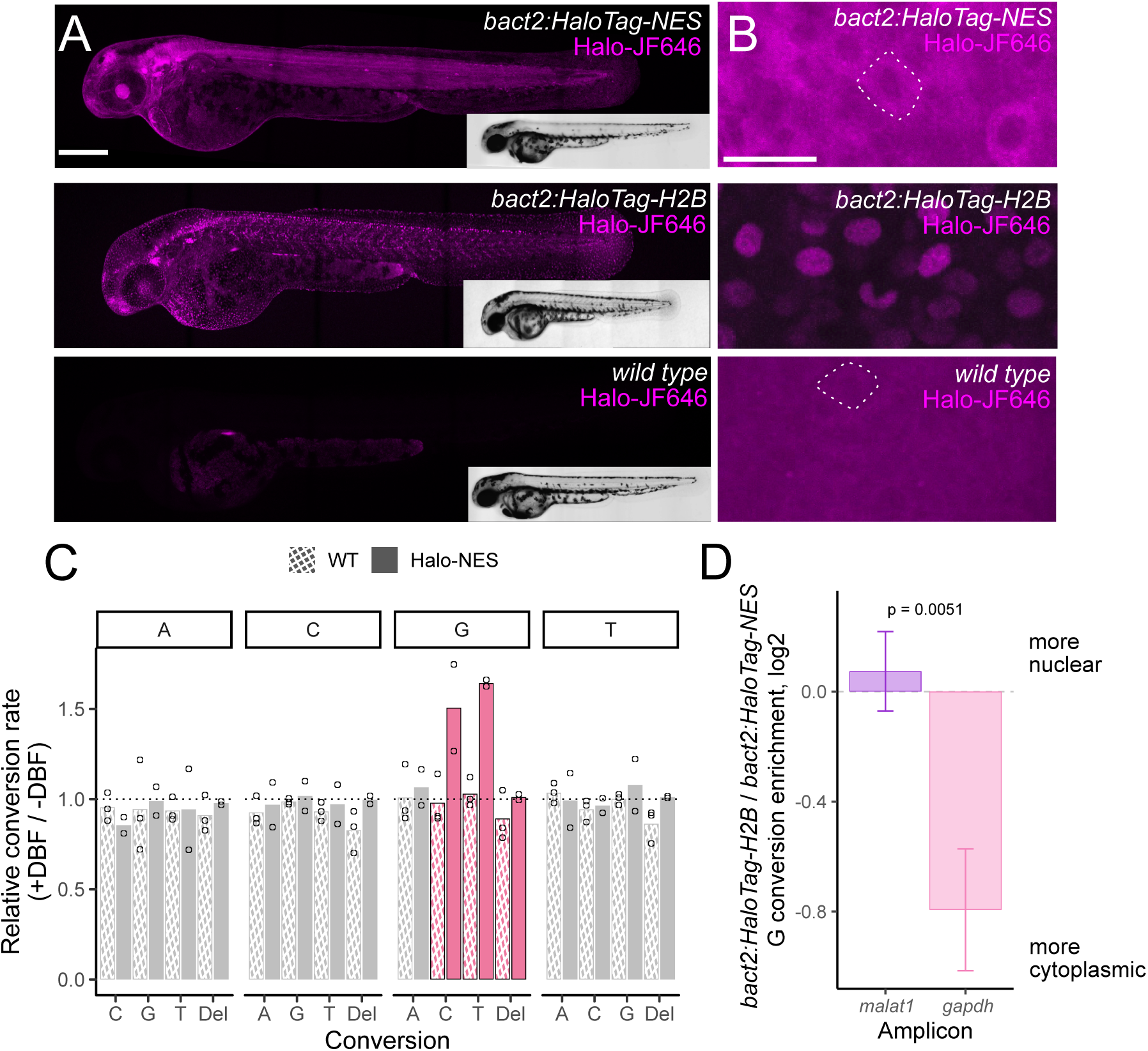
OINC-seq is compatible with RNA labeling in live animals. (A) Representative images of 2 days post-fertilization (dpf) homozygous bact2:Halo-NES transgenic zebrafish (top), bact2:Halo-H2B transgenic zebrafish (middle) or wildtype zebrafish embryo (bottom) stained with the fluorescent Halo ligand JF646 (magenta). Anterior to the left, scale bar 200 μm. (B) Confocal imaging of epithelial cells in homozygous bact2:Halo-NES (top), bact2:Halo-H2B (middle) or non-transgenic wild type (bottom). A single cell is highlighted by outlines. The HaloTag-NES fusion, visualized using JF646 (magenta), is excluded from the nucleus while the HaloTag-H2B fusion is localized to the nucleus. (C) Bulk relative conversion rates comparing Halo-DBF-treated and untreated samples for wild type (crosshatch, n=3) and bact2:Halo-NES (solid, n=2) zebrafish following OINC-seq at 2 dpf. (D) Relative enrichment of G conversions in malat1 and gapdh RT-PCR amplicons in HaloTag-H2B samples compared to HaloTag-NES samples.

We next assayed the ability of OINC-seq to create and detect RNA oxidation events in live zebrafish embryos. We incubated homozygous *bact2:HaloTag-NES* and wild type embryos at 2 dpf with Halo-DBF in the E3 embryo medium for 1 hour and then washed out unbound ligand for 2 hours (**Figure S5F**, see **Methods** for details). As a control, in a separate cohort of embryos, we omitted the Halo-DBF treatment. We then irradiated embryos in E3 medium with green light for 5 minutes and collected total RNA. We analyzed the resulting RNA samples using 3′ end RNAseq libraries and quantified oxidation-induced nucleotide conversions using PIGPEN.

Compared to zebrafish samples that were not treated with Halo-DBF, we observed a specific increase in G conversions in the Halo-DBF-treated samples, again consistent with OINC-seq inducing guanosine oxidation events (**Figure 5C**). The rates of G conversions, however, were markedly lower than observed in HeLa cells (**Figure 4**). This may be due to a lower expression of the transgene in zebrafish compared to HeLa cells, the transgene not being expressed in every cell in the embryo, or issues with penetration of the Halo-DBF ligand through zebrafish tissue. Conversely, G conversions were not induced by Halo-DBF treatment in wild type embryos, indicating that they required expression of the Halo fusion protein (**Figure 5C**).

To analyze the spatial specificity of labeling in zebrafish, we analyzed RT-PCR amplicons of *gapdh* and *malat1* transcripts from both *bact2:HaloTag-H2B* and *bact2:HaloTag-NES* lines as we had done previously with HeLa cell RNA (**Figure 3**). As before, for both transcripts, we observed more G conversions in Halo-DBF treated zebrafish than in untreated zebrafish, indicative of induced RNA oxidation (**Figure S5G-H**). The conversion rates of other nucleotides were relatively less affected, again showing the specificity of the reaction. The rates of G conversions, though, were again lower than we had observed in HeLa cells.

Still, to assess the spatial specificity of OINC-seq in zebrafish, we compared G conversion rates for both amplicons between the *bact2:HaloTag-H2B* and *bact2:HaloTag-NES* lines. We found that G conversions in *malat1* were relatively more enriched in the *bact2:HaloTag-H2B* line while *gapdh* conversions were relatively more enriched in the *bact2:HaloTag-NES* line (**Figure 5D**). These results were highly similar to those observed with the human orthologs of these RNAs in HeLa cells that had undergone nuclear or cytoplasmic labeling (**Figure 3**) and suggest spatial specific labeling in zebrafish. Taken together, these proof-of-principle experiments using transgenic HaloTag with nuclear exclusion sequence establish the functionality of OINC-seq *in vivo* using zebrafish embryos.

## DISCUSSION

A variety of proximity-based methods for RNA labeling and analysis have recently been developed (21–24) to enable the study of RNA localization at previously intractable subcellular locations. All of these approaches involve labeling RNA at a subcellular location and subsequently separating the labeled RNA from unlabeled RNA using biotin-mediated enrichment. Here, we present OINC-seq, an *in vivo*-compatible RNA proximity labeling method in which the *in situ*-deposited label is read out directly using high-throughput sequencing.

OINC-seq has several advantages over previously reported methods. First, removing the need for biotinylation and enrichment greatly streamlines the process. With OINC-seq, all steps necessary to create RNA samples ready for RNAseq library preparation can be performed in 2 hours. The same process can take up to 12 hours with earlier approaches. Second, removing the enrichment step removes a potential source of experimental noise. Streptavidin beads, although mostly inert, can nonspecifically bind unbiotinylated RNA (39). If the relative amount of biotinylated and unbiotinylated RNA varies across samples, the contribution of nonspecifically bound RNA can also vary, skewing results.

The ability of OINC-seq to detect multiple labeling events within a single RNA molecule greatly increases the discriminatory ability of OINC-seq to identify true, proximity-dependent labeling events. RNAs labeled in a proximity-dependent manner are more likely to have multiple labels across their length. Conversely, background labels introduced during the experiment in a nonspecific way are expected to be less abundant and occur more randomly across the RNA sample with each RNA molecule receiving relatively fewer labels. Because of the high-affinity binding between biotin and streptavidin, RNAs with a single biotin moiety and those with multiple biotin moieties may interact with streptavidin with similar efficiencies. Therefore, with biotin-based approaches, truly labeled RNAs and nonspecifically labeled RNAs may be enriched with similar efficiencies. In contrast, OINC-seq’s ability to quantify labeled residues rather than simply detect their presence or absence allows further filtering for transcripts with multiple labels. In fact, we observed that requiring multiple labels on a single RNA for quantification increased the ability of OINC-seq to discriminate between labeled and unlabeled RNA samples by 10-fold.

Finally, the majority of RNA localization studies have been conducted in cultured cell models. To better understand the effects of RNA localization in living organisms, labeling RNAs *in vivo* is desirable. However, biotinylation of RNAs *in vivo* often involves toxic reagents, technical challenges, and high costs. By circumventing biotinylation, OINC-seq enables labeling RNA in living organisms, as shown here in zebrafish embryos (**Figure 5**). The ubiquitous, inducible, tissue-specific expression of transgene-encoded HaloTag protein fusions with specific localization tags is applicable to numerous model organisms, expanding the opportunities to discover and quantify RNA localization *in vivo*. Still, RNA labeling in zebrafish embryos may require additional optimization to maximize rates of G conversion.

On the other hand, removing the enrichment step also has disadvantages. RNA oxidation events are quite rare, even in samples where the parameters that control the extent of proximity-induced RNA oxidation have been maximized. In these samples, between 1 in 1000 and 1 in 10,000 guanosine residues appear as cytidine or thymidine in the resulting cDNA, indicating their potential oxidation. In experiments where the fraction of RNA expected to be labeled is lower (e.g., when targeting a small subcellular region), the ability to detect a reasonable number of oxidation events at a reasonable read depth may become an issue. Biotinylation-based methods circumvent this problem by enriching these rare events. With OINC-seq, no such enrichment is performed.

RNA oxidation occurs in other contexts besides engineered proximity labeling experiments. RNAs are oxidized naturally in cells, such as a result of exposure to reactive oxygen species and/or carcinogens (32, 40). Mechanistically, the occurrence of 8OG in coding regions leads to ribosome stalling and mRNA decay through translation surveillance pathways (41). 8OG residues also inhibit RNA degradation by Xrn1 (42), possibly leading to a buildup of partially degraded transcripts. Increased levels of RNA oxidation is associated with a range of diseases, from neurodegenerative diseases to cancer (40). For these reasons, the detection of oxidized RNA in RNAseq samples may be useful beyond localization studies. PIGPEN can identify oxidized nucleotides in these samples, as evidenced by its ability to distinguish input and eluate samples from an 8OG immunoprecipitation experiment (**Figure S2B,C**). Further, since oxidation affects both RNA and DNA guanosine residues, care is needed to discriminate the effect of oxidation on RNA from DNA. By masking mutated positions (i.e. SNPs), PIGPEN is capable of identifying RNA-specific oxidation events. Finally, although here we paired PIGPEN with RNA oxidation induced by localized Halo-DBF, other proximity labeling approaches, including APEX-seq and CAP-seq, also utilize RNA oxidation. PIGPEN may therefore be useful in detecting and quantifying labeled RNA using variations of those techniques in which biotinylation is omitted.

We currently cannot explain with certainty why detected nucleotide conversion events are much higher in single transcript amplicon experiments (i.e. those involving *GAPDH* and *MALAT1*, ∼100 fold) than in transcriptome-scale experiments (∼10 fold). Because they only interrogate a single RNA species, amplicon experiments have many more reads per transcript species than transcriptome-scale experiments. However, this is not the reason for the difference in nucleotide conversion enrichments as subsampling the amplicon data did not change conversion enrichment values (**Figure S4A**). Instead, differences in sequencing library preparation approaches may contribute to this effect. Nucleotide conversion frequencies in control, untreated RNA were highly dependent upon library preparation method with transcriptome-wide libraries showing higher rates of “background” mutations in unlabeled RNA samples (**Figure S4B**). Consequently, the choice of library preparation strategy can have an effect on observed nucleotide conversion rates, likely through differences in fidelity in the polymerases used. Further, quantifying the localization of single transcripts, including reporter transcripts, using amplicons has numerous applications in developmental biology, disease modeling, drug discovery, and more.

Finally, given that OINC-seq is capable of inducing RNA labeling *in vivo*, OINC-seq has potential application for cell type-specific transcriptome analysis from bulk RNA-sequencing. After driving the transgenic expression of a non-localized HaloTag fusion protein with a tissue-specific promoter/enhancer, RNAs in the HaloTag-expressing cells would be specifically labeled, facilitating their discrimination from unlabeled RNAs in other tissues in bulk sequencing data. Similar tissue-or cell type-specific sequencing currently requires flow cytometry-based sorting of cells of interest followed by RNA sequencing. OINC-seq may therefore offer a streamlined experimental workflow for tissue-specific transcriptomics applications in model organisms.

## ACKNOWLEDGEMENTS

We thank members of the Taliaferro lab for helpful discussions regarding experiments and analyses, and Rob Patro for helpful discussions and software regarding RNA oxidation quantification. We also thank Harrison Wells and Dr. Robert Lalonde for assistance with zebrafish imaging. Portions of the figures in this work were created by BioRender. This work was supported by NIH grants R35-GM133385 (J.M.T.), R35-GM140813 (C.G.P.), T32-GM008730 (H.Y.G.L. and R.G.), T32-CA190216 (A.M.), and 2R15-GM132816-02 (M.J.E.R.). It was also supported by the W.M. Keck Foundation (J.M.T. and C.G.P.), the RNA Bioscience Initiative at the University of Colorado Anschutz Medical Campus (H.Y.G.L., R.G., L.K.W. and J.M.T.), the Swiss National Science Foundation (Sinergia grant CRSII5_180345, C.M.), the Swiss Bridge Foundation (C.M.); the University of Colorado School of Medicine Anschutz Medical Campus and the Children’s Hospital Colorado Foundation (C.M.); C.M. holds the Helen and Arthur E. Johnson Chair for the Cardiac Research Director. We also thank Lily Nguyen for her unwavering support.

## DATA AVAILABILITY

High-throughput sequencing data associated with this manuscript has been deposited at the Gene Expression Omnibus under accession number GSE279714.

## DECLARATION OF INTERESTS

The authors declare no competing interests.

## MATERIALS AND METHODS

### Creation of stably-integrated transgenic cell lines expressing HaloTag-fusion proteins

All transgenic cell lines started as HeLa cell lines containing a single *loxP* cassette with blasticidin resistance (43). These HeLa cells were co-transfected with 1-2 µg of a plasmid containing the HaloTag fusion and puromycin resistance, alongside 50-100 ng of a plasmid that expressed *Cre* recombinase (Addgene pBT140). Transfection was performed with Lipofectamine 2000 (Thermo Scientific) in accordance with the manufacturer’s instructions. Cells were allowed to recover for 48 h post transfection prior to selection with puromycin at 5 µg/mL. A stable population of integrants derived from numerous selected colonies were pooled to reduce variability from single clones.

Expression of the HaloTag fusion gene product is controlled by a reverse tetracycline-controlled transactivator (rtTA) and is thus doxycycline-dependent. To express the HaloTag fusion, cells were incubated with 1 µg/mL of doxycycline for 48 h prior to experiments. All Halo-Tag fusion protein constructs contained N-terminally tagged HaloTags (Halo-p65, Halo-P450, Halo-ATP5MC1, and Halo-MAVS) with the exception of Halo-H2B, which contained a C-terminally tagged HaloTag.

### Validation of subcellular localization of HaloTag fusion proteins

OINC-seq requires localization of HaloTag fusion proteins to the subcellular location of interest. To verify proper localization of HaloTag fusions in each cell line, cells expressing a Halo-tagged protein were plated in 12-well plates on poly-D-lysine-coated coverslips (Neuvitro cat #H-22-15-PDL). Doxycycline at 1 µg/mL was added to induce expression of the HaloTag fusion for 48 h. On the day of the experiment, the media was removed and the cells washed in PBS once. Cells were then incubated with a fluorescent Halo ligand (Janelia Fluor 594) or Oregon Green (Promega) at 25 nM in PBS for 15 min. Excess ligand was washed off with 3 washes in complete media (DMEM supplemented with penicillin, streptomycin and 10% Equafetal (Atlas Biosciences)) in the tissue culture hood for a total of 30 min. Cells were then fixed in 3.7% formaldehyde (10% formalin) for 12 min at room temperature (RT). Cells were then washed with PBS. For experiments requiring co-localization with primary antibodies recognizing subcellular structures, cells were blocked with 5% BSA for 1 h followed by primary antibody staining overnight at 4°C in 3% BSA. Primary antibodies and dilutions used: rabbit polyclonal GRP450 used at 1:500 (gift from Christopher Nicchitta) and rabbit polyclonal TOM20 at 1:250 (Proteintech cat. #11802-1-AP, lot #96707). Cells were then washed in 0.005% Tween in PBS (PBST) and stained with Alexafluor anti-Rabbit IgG 555 (Cell Signaling cat. #4414S, lot #16) for 1 h at RT. Excess secondary antibodies were washed off with PBST. DAPI was added at 100 ng/mL for 10 min. Cells were washed again with PBS and coverslips were mounted and imaged.

Subcellular visualization of the inner and outer mitochondrial proteins required amendments to the procedure described above. MitoView Green (Biotium cat. #70054, lot #16M0316-1107140) staining was performed in live cells following incubation of the fluorescent Halo ligand for 30 min in PBS. Excess MitoView was washed off with complete media DMEM supplemented with penicillin, streptomycin and 10% Equafetal (Atlas Biosciences) for 15 min. Cells were then fixed, treated with DAPI, and mounted as described previously. Imaging was performed using structured illumination microscopy (SIM) using a Nikon SIM (N-SIM) with a Nikon Ti2 (Nikon Instruments; LU-N3-SIM) microscope equipped with a 100× SR Apo TIRF, NA 1.49 objective. Images were captured using a Hamamatsu ORCA-Flash 4.0 Digital CMOS camera (C13440) with 0.1-µm Z steps. All images were collected at 25°C using NIS Elements software (Nikon). Raw SIM images were reconstructed using the image slice reconstruction algorithm (NIS Elements). Line scan analyses were conducted on 100 cells per labeling condition and alignment was determined by the brightest pixel intensity in the MitoView image channel. Line scans represent the average of all measured cells at each position.

### In-cell visualization of 8OG-containing RNA

Cells expressing the Halo-p65 fusion protein were grown in 12-well plates on poly-D-lysine-coated coverslips. Expression of the HaloTag-fusion protein was induced as previously described. On the day of the experiment, complete media was removed and cells were incubated with PBS either including (+Halo-DBF) or lacking (-Halo-DBF) 5 µM Halo-DBF for 15 min. Excess Halo-DBF was removed with 3 washes in complete media in the tissue culture hood for a total of 30 min. Cells were washed in PBS prior to the labeling. The dishes of cells were then sandwiched between two green LED flood lights (AC: 85-265V, light source: 144pcs SMD2835 LED, power: 100W; manufacturer: T-SUNRISE, cat. #B01N1S6D8K) in a dark room as previously described (25). All cells are then subjected to green light exposure for a predetermined period of time, typically 5 minutes. Cells were immediately fixed with 100%, pre-chilled methanol at −20°C for 10 min. Fixed cells were washed 3 times with PBS for a total of 15 min. Fixed cells were cleared with 0.005% PBST for 1 h and blocked with 5% BSA for 1 h at RT. Cells were incubated with a mouse antibody recognizing 8-oxoguanosine (Santa Cruz, cat. #sc-66046, lot # L0112) at 1:500 at 4°C overnight in 3% BSA. Excess primary antibody was washed off with PBST. Secondary antibody staining was performed with Alexafluor anti-mouse IgG 488 (Cell Signaling cat. #4408S, lot #17) in PBST for 1 h at RT. Excess secondary antibody was washed off and DAPI was added at 100 ng/mL for 10 min. Coverslips were then mounted and imaged. The same exposure time and laser power was used for both the +/− Halo-DBF conditions and all further image processing was performed in FIJI (44) with identical parameters.

### Synthesis of RNA containing the Guanidinohydantoin (Gh) Lesion

Obtained using an adapted procedure from previous reports (29). A solution of RNA (6 nmol), containing 8-oxoguanosine (42), in RNase-free H_2_O (50 µL was incubated at 0°C (10 minutes); followed by addition of a Na_2_IrCl_6_ aqueous solution (16 µL, 4.5 mM, 72 nmol) and further incubation for 30 min. Purification was then carried out using HPLC on a standard C18 column: solvent A = 0.1M TEAA (pH=7) & solvent B = 1:1 ACN/ 0.1M TEAA with a gradient from 15% B to 100% B over 45 min at a flow rate of 1mL/min while monitoring absorbance at 260 nm. The peak of interest was collected (27 min), concentrated under reduced pressure to yield the RNA containing the Gh lesion. The oligonucleotide was characterized via MALDI-TOF, as previously described (45).

### Generation of oligonucleotides containing 8OG or GH

We generated synthetic oligonucleotide substrates with an 8OG, GH, or unmodified guanosine at a known nucleotide position via a splint ligation strategy (**Figure S1C**). A 29 nt RNA oligo containing an 8OG at the 12th position was synthesized as previously described (42). The same RNA oligo containing a GH at the same position was generated as described above. An unmodified oligonucleotide of identical sequence was ordered from Integrated DNA Technologies. Additional sequence containing PCR handles was added to each oligo via ligation of flanking RNA oligonucleotides (**Figure S1C**). The 89 nt product was phosphorylated with T4 PNK and a final 20 nt RNA oligo with sequence GGCUUCGCAGUCCUUAGAAG (Chemgenes) was ligated to the 5’ end of the 89 mer using T4 RNA Ligase 1 (NEB). Finally, the product was gel purified to isolate the desired 109 nt product.

### Analysis of 8OG or GH oligo sequencing errors

Libraries were prepared with RNA oligos containing either a normal Guanosine (control) or an oxidized lesion (8OG or GH) at a known position. Primers used can be found in **Supplemental Table 1**. Libraries were sequenced using paired end sequencing (2 x 150bp) on a NovaSeq platform (Illumina). 2-5 million read pairs were sequenced for each replicate. Only the forward reads were used for downstream analyses. The sequence of the oligos of interest were pulled from raw sequencing reads using Python’s Regular Expression Syntax (REGEX) allowing for up to 4 mismatches. Each filtered read was queried against the control sequence and mismatches relative to the control sequence were quantified.

### OINC-Seq labeling and RNA isolation

Cells expressing a single HaloTag fusion protein were grown in either 6-well plates or 10 cm dishes (1 well or dish per replicate per condition) and expression of the HaloTag fusion protein was induced with doxycycline. On the day of the experiment, cells were 80-90% confluent. Cells were then incubated with Halo-DBF (5 µM, unless otherwise specified, in PBS) at 37°C for 15 min (+Halo-DBF). Control cells did not receive Halo-DBF treatment (-Halo-DBF). Excess Halo-DBF was washed off with complete media 3 times for a total of 30 min. Media was then replaced with PBS and cells were irradiated with green light from an LED panel (5 min, unless otherwise specified) as previously described (25).

Total RNA was then immediately isolated from the cells with Trizol (Ambion) following manufacturer’s instructions with an additional step of trituration through a 20G needle 20 times. Contaminating DNA was removed by incubating the purified RNA with DNase I (Thermo Scientific) for 30 min at 37°C followed by RNA recovery with an RNA recovery kit (Zymogen Quick-RNA MicroPrep, cat. #R1051). RNA quality and concentrations were checked via Nanodrop prior to downstream applications. All samples were required to have a 260/280 > 2.0, a 260/230 > 2.0, and an RNA concentration of ≥ 500 ng/µL.

### OINC-Seq library preparation and high-throughput sequencing

For single amplicon libraries, 5 µg of RNA was reverse transcribed with SuperScript IV (ThermoFisher cat #18090050) following the manufacturer’s protocol. Following cDNA synthesis, RNA was degraded with 5 units of each RNase H (BioLabs cat #M029L) and RNase A/T1 (Thermo Scientific cat #EN0551). Library amplification was performed using Q5 polymerase (NEB 0492) with primers that contain Illumina handles and unique barcodes for demultiplexing of replicates and conditions (sequences in **Supplemental Table 1**). Annealing temperatures and PCR cycle counts were determined experimentally for each gene (**Supplemental Table 1**). Libraries were isolated using a DNA isolation kit (Zymogen DNA Clean & Concentrator cat #D4014). Primer dimers were removed with a 0.75x magnetic bead cleanup (AxePrep cat #MAG-PCR-CL-5). Sample concentrations were determined by Qubit Fluorometer and quality check performed by Tapestation (Agilent High Sensitivity DNA ScreenTape #5067-5584).

Transcriptome-wide libraries were generated from 200 ng of RNA using the Lexogen’s QuantSeq 3’ mRNA-Seq kit (Lexogen cat #015) following the manufacturer’s protocol. Depending on the input, 14-20 PCR cycles were used to amplify libraries.

Libraries were sequenced using paired end sequencing (2 x 150bp) on a NovaSeq platform (Illumina). Read depth varied depending on experiments. Typically, 1-3 million read pairs were sequenced for each replicate for single library amplicons, approximately 50 million read pairs were sequenced for each replicon for transcriptome-wide studies, and 100 million read pairs were sequenced for each replicon for the *in vivo* zebrafish studies.

### Adapter trimming of 3’ end sequencing libraries

For transcriptome-wide OINC-seq experiments, 3’ end RNAseq libraries were prepared (Lexogen) and sequenced using paired-end sequencing. For these samples, adapters were removed from reads using Cutadapt (46) and the following strategy:

Step 1. The UMI (the first 6 nt of read 1) was removed and the 3’ adapter (AAAAAAAAAAAAAAAAAAAA) was trimmed off of read 1.

Step 2. The 5’ adapter (TTTTTTTTTTTTTTTTTTTT) was trimmed from read 2.

Step 3. A 3’ adapter (AGATCGGAAGAGCGTCGTGTAGGGAAAGACGGTA) was attempted to be trimmed off of read 2. If reads did not contain this adapter, their trimming was completed and they were saved. If reads did contain this adapter, they were further processed by step 4.

Step 4. The last 6 bases (which correspond to the UMI) were removed from read 2. Step 5. Trimmed reads from step 4 were combined with untrimmed reads from step 3.

### Analysis of OINC-seq data with PIGPEN

Nucleotide conversion frequencies in OINC-seq data were derived using PIGPEN software (https://github.com/ <u>TaliaferroLab/OINC-seq</u>). In brief, PIGPEN takes in RNAseq reads and aligns them to a reference genome using STAR (47). These alignments are used to identify and quantify nucleotide conversions relative to the reference genome. If desired, genome positions with high levels of mutations (e.g. SNPs) can be called using varscan (48) and masked from further analysis. As a default, in order for a site to be masked a nucleotide must be covered by at least 20 reads and at least 20% of those reads must contain a SNP at that nucleotide. Low quality score nucleotides are masked (default Q < 30), and, if desired, a requirement that a given conversion be observed in both mates of a read pair can also be enforced.

In parallel, reads are assigned to transcripts using salmon (30), and the fractional assignment of each read to a transcript is derived using postmaster (https://github.com/COMBINE-lab/postmaster). At the end of this process, each read has the number of conversions it contains quantified and has also been fractionally assigned to one or more transcripts. For each read, these two measurements are combined to assign nucleotide conversions to transcripts. For each gene, nucleotide conversions are then summed across all substituent transcripts to give gene-level conversion counts.

For the transcriptome-scale experiments, ER-localized RNAs were defined as those previously described as “high-confidence” ER-localized RNAs in a previous study (35). RNAs defined as those localized to the outer mitochondrial membrane were those encoding proteins with the gene ontology term “Mitochondrial Protein Complex” (GO:0098798). RNAs defined as those localized to the mitochondrial matrix were those that originated from genes on the mitochondrial chromosome.

To identify genes whose G to T and G to C conversion rates change across samples or conditions, the Bioinformatic Analysis of the Conversion Of Nucleotides (BACON) module of PIGPEN was used (<u>https://</u> <u>github.com/TaliaferroLab/OINC-seq</u>). BACON fits a binomial generalized linear mixed effects model of converted G counts and nonconverted G counts against conditions. This model is then compared to a null model in which the effect of the condition was removed. A likelihood ratio test was then used to evaluate the relative fit between the experimental and null models. P values were derived from the likelihood ratio test and then corrected for multiple hypothesis testing using a Benjamini-Hochberg correction (49).

### Zebrafish husbandry and procedures

Animal care and procedures were carried out in accordance with the IACUC of the University of Colorado School of Medicine (protocol 00979), Aurora, Colorado, USA. All zebrafish embryos were incubated at 28.5°C in E3 medium unless indicated otherwise. Staging was performed as per standard staging series (50).

### Zebrafish Expression constructs

Gateway 5’ entry vector (*pE5’*) *bactin2* contains a 5.3-kb promoter element from the β-actin gene which drives expression broadly throughout the embryo (51, 52) (a gift from Dr. Kristin Artinger). The Gateway middle entry vector *pME_HaloTag-NES* was designed to contain the coding sequences for a 3xHA tag, followed by the Halotag, 3x MAPKK2 NES, P2A, mCerulean-CAAX sequences for membrane localization. The *HaloTag-NES* ORF was synthesized by Twist Biosciences and cloned into the *pENTR/D-TOPO* vector (Invitrogen cat #K240020) according to the manufacturer’s protocol.

Multisite Gateway recombination to generate the transgene expression clone was performed as described in the Invitrogen Multisite Gateway Manual (Invitrogen), with minor modifications. 10 fmol of *p5E_bactin2*, *pME_HaloTag-NES*, and *p3E_ubb* vectors were combined with 20 fmol of *pDestTol2* destination vector (*Tol2 kit #394*) (52), 1 µl of LR Clonase II Plus Enzyme Mix (Invitrogen, 12538120; vortexed twice for 2 s prior to use) and deionised H2O for a final reaction volume of 5 µl. Vector calculations for molarity were performed using the Multisite Gateway Excel spreadsheet (53). Reactions were incubated at 25°C overnight and treated the following day with 1 µl of 2 µg/µl Proteinase K for 15 min at 37°C. 3 µl of the reaction was transformed with One Shot TOP10 Chemically Competent E. coli (Invitrogen cat #C404010) and plated on Ampicillin selection plates. 5 colonies were cultured overnight in Ampicillin LB at 37°C. The cultures were miniprepped with the ZymoPURE plasmid miniprep kit (Zymogen cat #D4212) and plasmid integrity and correct gateway assembly was verified with restriction digests (BamHI, XhoI) and Sanger sequencing. One correct plasmid was chosen for further experiments and the transgenic zebrafish line creation.

### Generation of HaloTag transgenic zebrafish lines

Tol2 transposase-encoding capped mRNA was created by *in vitro* transcription using the SP6 mMessage mMachine Kit (Ambion, cat #AM1340) from the *pCS2+Tol2* plasmid (52) linearized by NotI restriction digest. RNA was purified with lithium chloride precipitation followed by the Megaclear Kit (Ambion cat #AM1908). 1 nL injection mix containing the *bact2:HaloTag-NES-2A-mCerulean-CAAX* plasmid DNA at 25 ng/µl and Tol2 mRNA at 25 ng/µl was injected into zebrafish AB wild-type embryos in the cell at the one-cell stage. The Tol2-based zebrafish transgenics were generated using standard experimental protocols as previously specified (54) to achieve single-insertion transgenics and reproducible quality control as follows. The injected embryos were raised to adulthood (F0 generation), and potential transgenic zebrafish founders were outcrossed with wild-type zebrafish to identify germline transmission. F1 progeny expressing ubiquitous mCerulean were selected and bred to establish stable transgenic zebrafish lines and F2 generation heterozygous carriers were selected for predicted Mendelian ratios of 50% transgene carriers in the offspring to establish a single-integration transgenic line. The heterozygous, transgenic F2 zebrafish were incrossed to yield homozygous F3 progeny that was then further incrossed as adults to retain homozygous breeding pairs to perform the experiments on.

### Zebrafish fluorophore treatment and imaging

Routine fluorescence observations were performed on a Leica M205FA dissecting microscope with a DFC450 C camera and 1.0x PlanApo M-Series objective, illuminated with a TL5000 light base and CoolLED pE-300white Illumination System.

The embryos of the established F3 transgenic zebrafish line *bact2:HaloTag-NES-2A-mCerulean-CAAX* (further denoted as HaloTag-NES) line were manually dechorionated at 2 dpf and treated with JF646 fluorophore (0.6 µM dissolved in E3) for 1 h at 28°C. Post-treatment, the zebrafish were washed in E3 (twice for 2 h in total) and then imaged. Prior to imaging, the zebrafish were anesthetized with 0.016% Tricaine-S (MS-222, Pentair Aquatic Ecosystems, Apopka, Florida, NC0342409) in E3 embryo medium and embedded in E3 with 1% low-melting-point agarose (Sigma Aldrich, A9045) on glass bottom culture dishes (Greiner Bio-One, Kremsmunster, Austria, 627861). Laser scanning confocal microscopy was performed on a Zeiss LSM880, with a 10x/1 air-objective lens for general overview and 20x/1.3 air-objective lens for zoom of the tail to capture the specific fluorescence signal from the fluorophore. Image analysis was performed with ImageJ/Fiji (44).

### Zebrafish OINC-Seq labeling

HaloTag-NES embryos at 2 dpf were dechorionated and treated with Halo-DBF (50µM dissolved in E3) for 1 h at 25°C (+Halo-DBF). Control wild type embryos did not receive this treatment (-Halo-DBF). To remove excess DBF, the embryos were washed in excess E3 (twice for 2 h in total). The zebrafish were transferred to a 6 cm dish and anesthetized with 0.016% Tricaine-S. The zebrafish were then labeled for 5 min with exposure to green light. RNA was immediately extracted from the zebrafish embryos with Trizol following manufacturer’s instructions with an additional step of trituration through a 20G needle 30 times. Samples were treated identically to all other OINC-Seq samples in further downstream applications, which have been described above.

## SUPPLEMENTARY TABLES

**Supplemental Table 1**: Primers used for library preparations. Primers include anchor sequences for illumina sequencing (gray), illumina i7 handles (dark blue), i5 handles (pink), internal barcodes (black), gene specific primers (light blue), and overlapping primer sequences (green). Corresponding species, gene, PCR cycle numbers and annealing temperatures are included.

**Supplemental Table 2**: BACON output comparing G conversion rates in OINC-seq samples labeled with p450 (ER localized) and p65 (cytoplasm localized). Genes with fewer than 100 reads in any sample are filtered out. Differences in G conversion rates are reported as ER - cytoplasm.

**Supplemental Table 3**: BACON output comparing G conversion rates in OINC-seq samples labeled with ATP5MC1 (mitochondria localized) and p65 (cytoplasm localized). Genes with fewer than 100 reads in any sample are filtered out. Differences in G conversion rates are reported as mitochondria - cytoplasm.

**Supplemental Table 4**: BACON output comparing G conversion rates in OINC-seq samples labeled with H2B (nucleus localized) and p65 (cytoplasm localized). Genes with fewer than 100 reads in any sample are filtered out. Differences in G conversion rates are reported as nucleus - cytoplasm.

## SUPPLEMENTARY FIGURES

**Figure S1:**
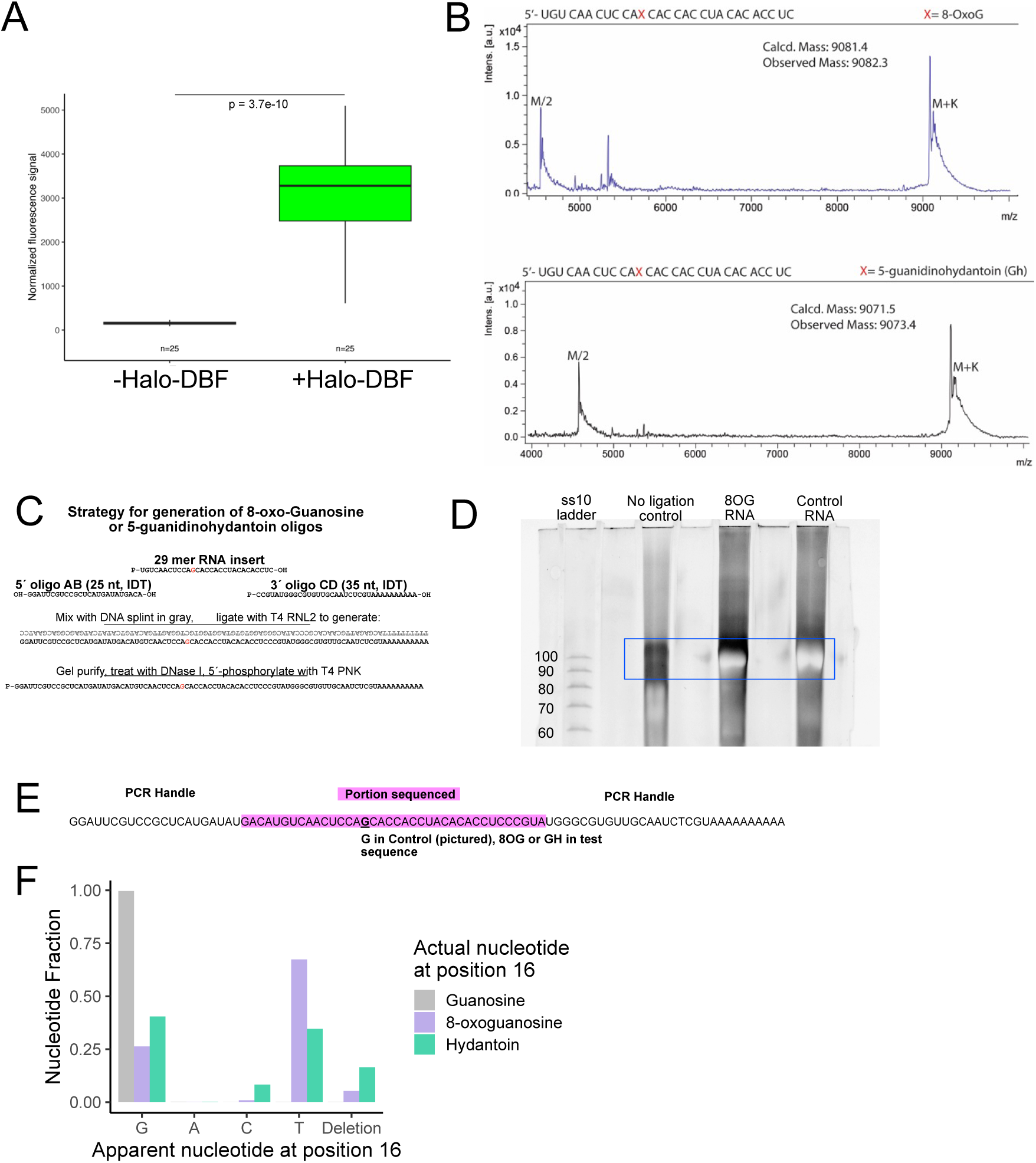
(A) Quantified normalized fluorescence of Halo-p65-expressing cells stained with an 8OG antibody that were treated or untreated with Halo-DBF. (B) MALDI-TOF corresponding to oligonucleotides of RNA containing 8-oxoG and Gh. (C) Schematic for synthesis of 8OG synthetic oligo products. (D) Polyacrylamide gel electrophoresis of ligated products. The lack of an inverted band in the control without the ligase and the presence of inverted bands in both the 8OG ligated product and the control ligated product at the correct size (109nt) is evidence of successful splinting of PCR handle onto synthesized RNA. (E) Synthesized RNA with PCR handles for OINC-seq. Portion sequenced by RNA-sequencing is highlighted in magenta. The underlined guanosine is either a normal guanosine (control) or a chemically synthesized 8-oxoguanosine (8OG) or 5-guanidinohydantoin (GH). (F)Bar graphs depicting the relative fractions of each nucleotide at position 16 in either the control sample with a G, or the oxidized samples with a 8OG or GH.

**Figure S2:**
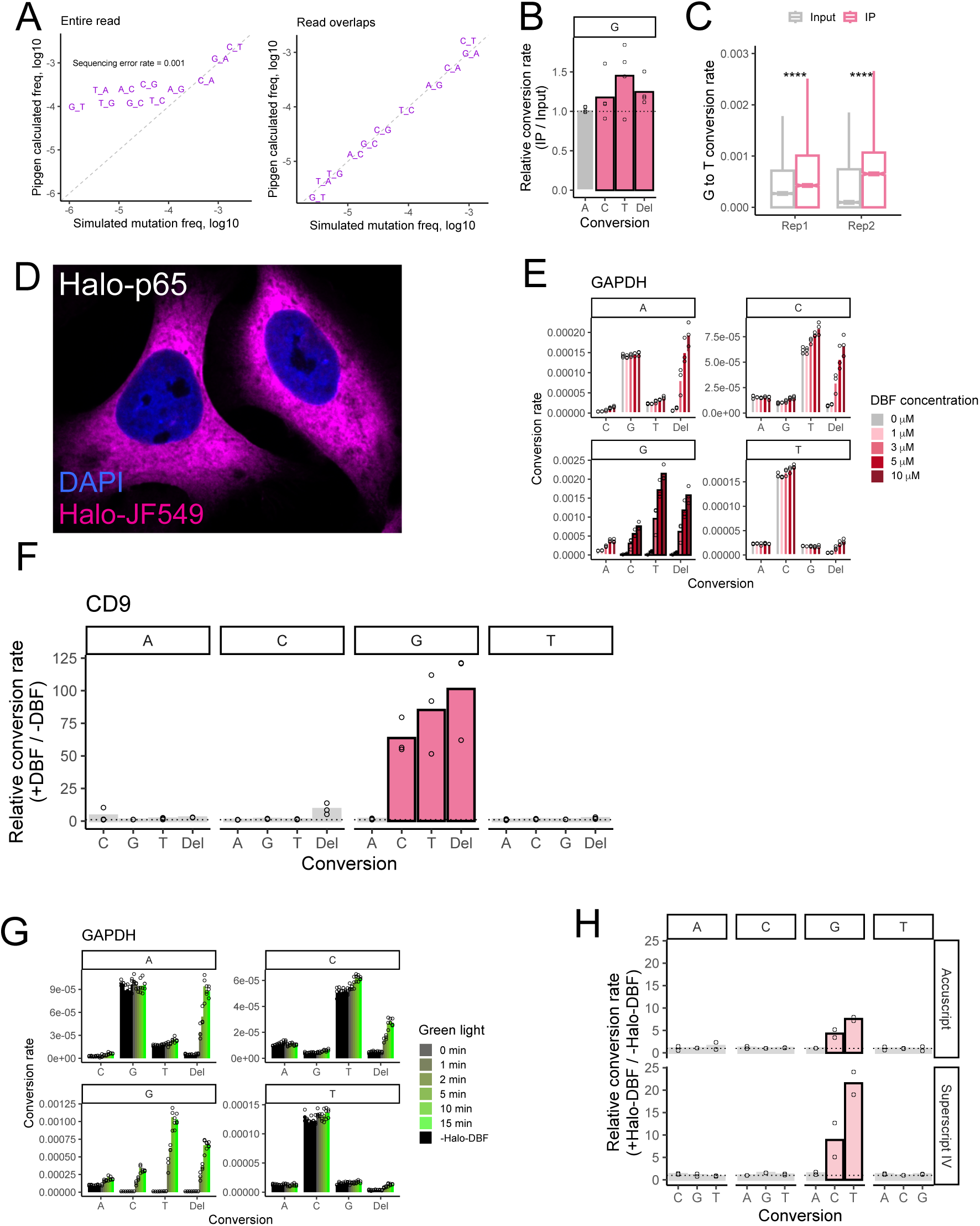
(A) Testing of PIGPEN using synthetic RNAseq data with defined nucleotide conversion rates. Both plots include a simulated sequencing error rate of 0.001. On the left, conversions were not required to be found in both reads of a mate pair, and sequencing error dominates. This effect is removed on the right when conversions were required to be found in both reads. (B) Bulk conversion rates comparing immunoprecipitated and input samples from a previously reported 8OG immunoprecipitation experiment. (C) G to T conversion rates across all genes of input and IP samples from a previously reported 8OG immunoprecipitation experiment. (D) Representative image of HeLa cells expressing the cytoplasmic HaloTag fusion p65-Halo. The Halo fusion is visualized with the fluorescent Halo ligand JF549 (magenta). Nuclei are stained with DAPI. (E) Rates of nucleotide conversions in the GAPDH RT-PCR amplicon as a function of increasing Halo-DBF concentration (n=3). In this analysis, conversions must be seen in both reads of a mate pair. (F) Quantification of conversion rates in an amplicon of the CD9 transcript in untreated cells and those treated with 5 µM Halo-DBF. (G) As in E, but plotting nucleotide conversion rates as a function of increasing exposure times to green light (n=3). Black bars represent samples in which Halo-DBF was omitted. (H) Relative nucleotide conversion rates between Halo-DBF treated and untreated samples in a Halo-p65 OINC-seq experiment in which either Accuscript or Superscript IV was used as the reverse transcriptase (n=2).

**Figure S3:**
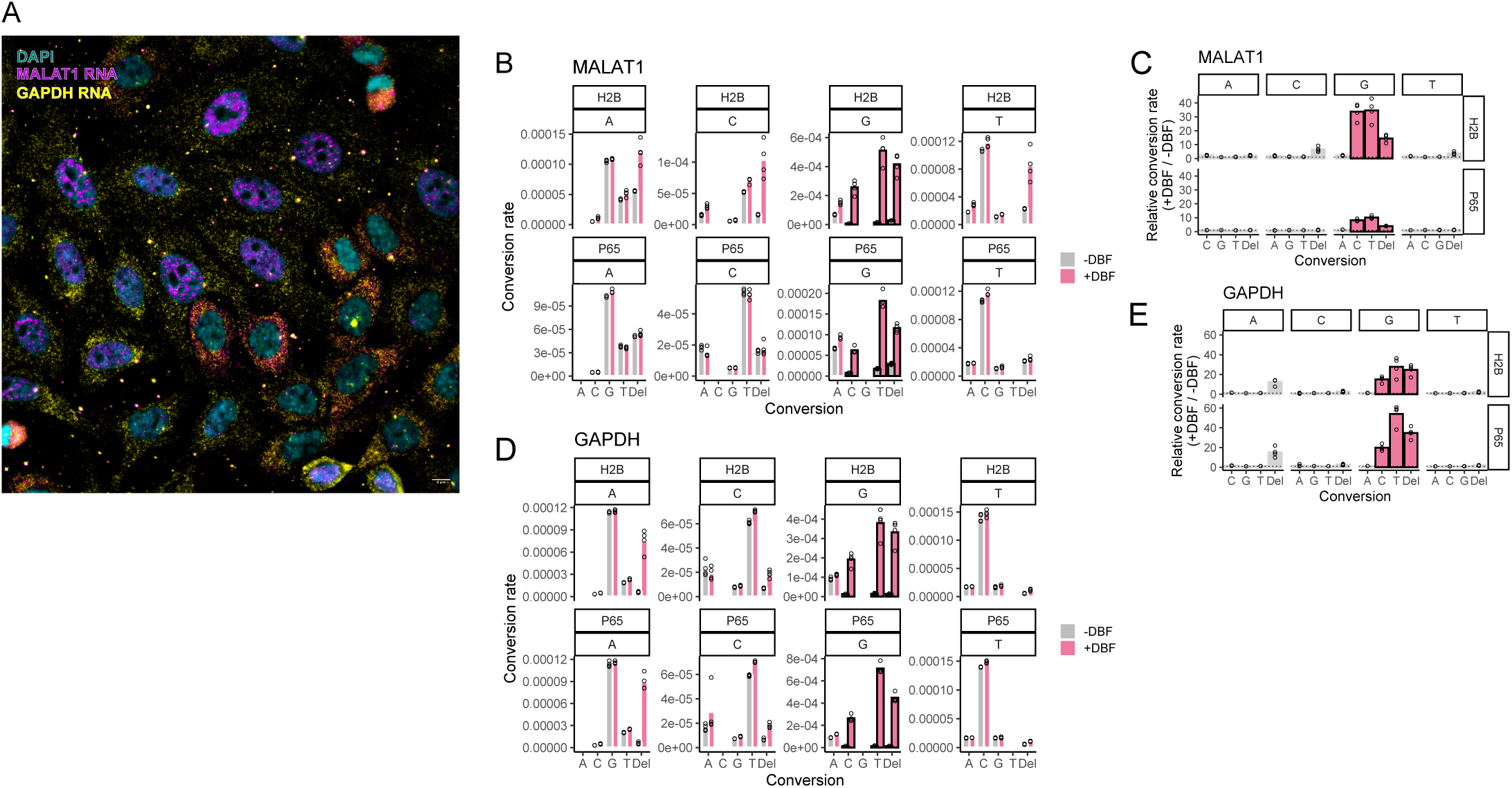
(A) smFISH imaging for MALAT1 RNA (magenta) and GAPDH RNA (yellow). DAPI is shown in cyan. Notice that while in most cells MALAT1 RNA is nuclear, in two cells near the center it is cytoplasmic. These cells likely recently underwent mitosis. The scalebar in the lower right corner is 8 microns. (B) Rates of nucleotide conversions in the MALAT1 RT-PCR amplicon in OINC-seq experiments in which the indicated Halo fusion (H2B or p65) was used. Analysis required observing conversions in both reads of a mate pair (n=3). (C) Relative nucleotide conversion rates in the MALAT1 amplicon between Halo-DBF treated and untreated samples in OINC-seq experiments in which the indicated Halo fusion (H2B or p65) was used (n=3). Analysis required observing conversions in both reads of a mate pair and one G conversion. (D) As in A, but for the GAPDH RT-PCR amplicon (n=3). E. As in B, but for the GAPDH RT-PCR amplicon (n=3).

**Figure S4:**
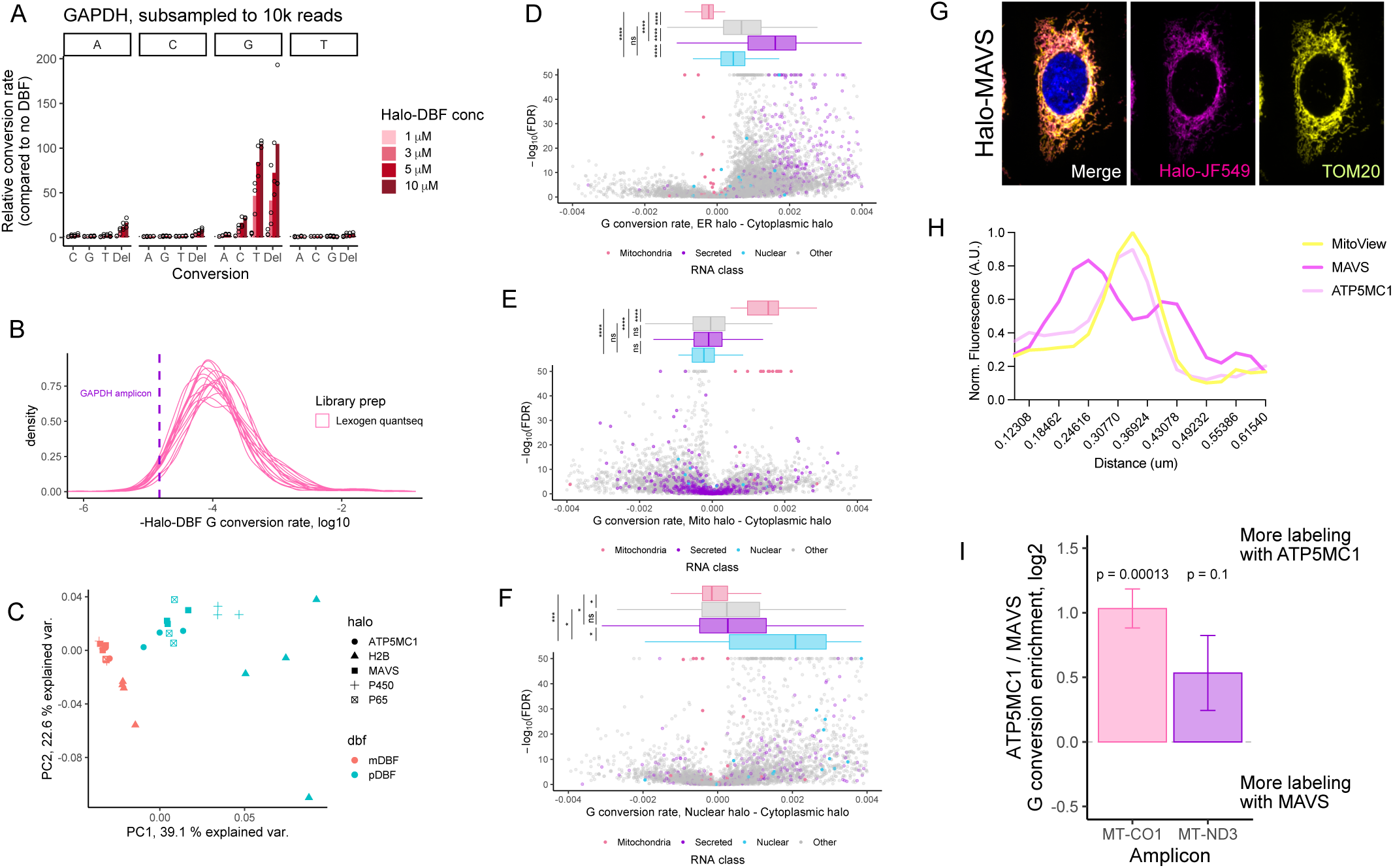
(A) As in figure 2C, proximity-mediated RNA oxidation was induced using a cytoplasmically-localized HaloTag protein and increasing amounts of the oxygen radical-producing Halo-DBF ligand. Relative conversion rates in an RT-PCR amplicon of the GAPDH transcript were calculated and compared to samples in which Halo-DBF was omitted (n=3). In this case, to assess the contribution of read depth to observed conversion rates, 10,000 reads were randomly sampled from the approximately one million obtained. For this analysis, conversions were required to be seen in both reads of a mate pair, and only one G to T or G to C conversion in a read was required. (B) G conversion rates in samples in which Halo-DBF was omitted (i.e. “background” conversion rates). Data from multiple transcriptome-wide, as well as the observed background G conversion rate in the GAPDH amplicon experiments (n=2). (C) PCA of G conversion rates for transcriptome-wide samples. (D) Volcano plots of genes comparing G conversion rates in the ER-targeted (p450) to broadly cytoplasmically targeted (p65) Halo fusions. P values and FDR values were calculated using BACON’s statistical framework. (E) As in D, but comparing mitochondrially targeted (ATP5MC1) to cytoplasmically targeted (p65) Halo fusions. (F) As in D, but comparing nuclearly targeted (H2B) to cytoplasmically targeted (p65) Halo fusions. (G) Fluorescence microscopy of the outer mitochondrial membrane HaloTag fusion (Halo-MAVS, magenta) co-stained with the mitochondrial marker TOM20 (yellow). (H) Line scan analyses quantifying co-localization of Halo-ATP5MC1 (light pink) or Halo-MAVS (magenta) with MitoView (yellow). Each line represents the average measurement across 100 cells. (I) Relative enrichment of G conversions in ATP5MC1 (mitochondrial matrix) and MAVS (outer mitochondrial membrane) samples for amplicons of two mitochondrial chromosome-encoded RNAs. P values were calculated using a T-test to ask if the mean of the values was different than 0.

**Figure S5:**
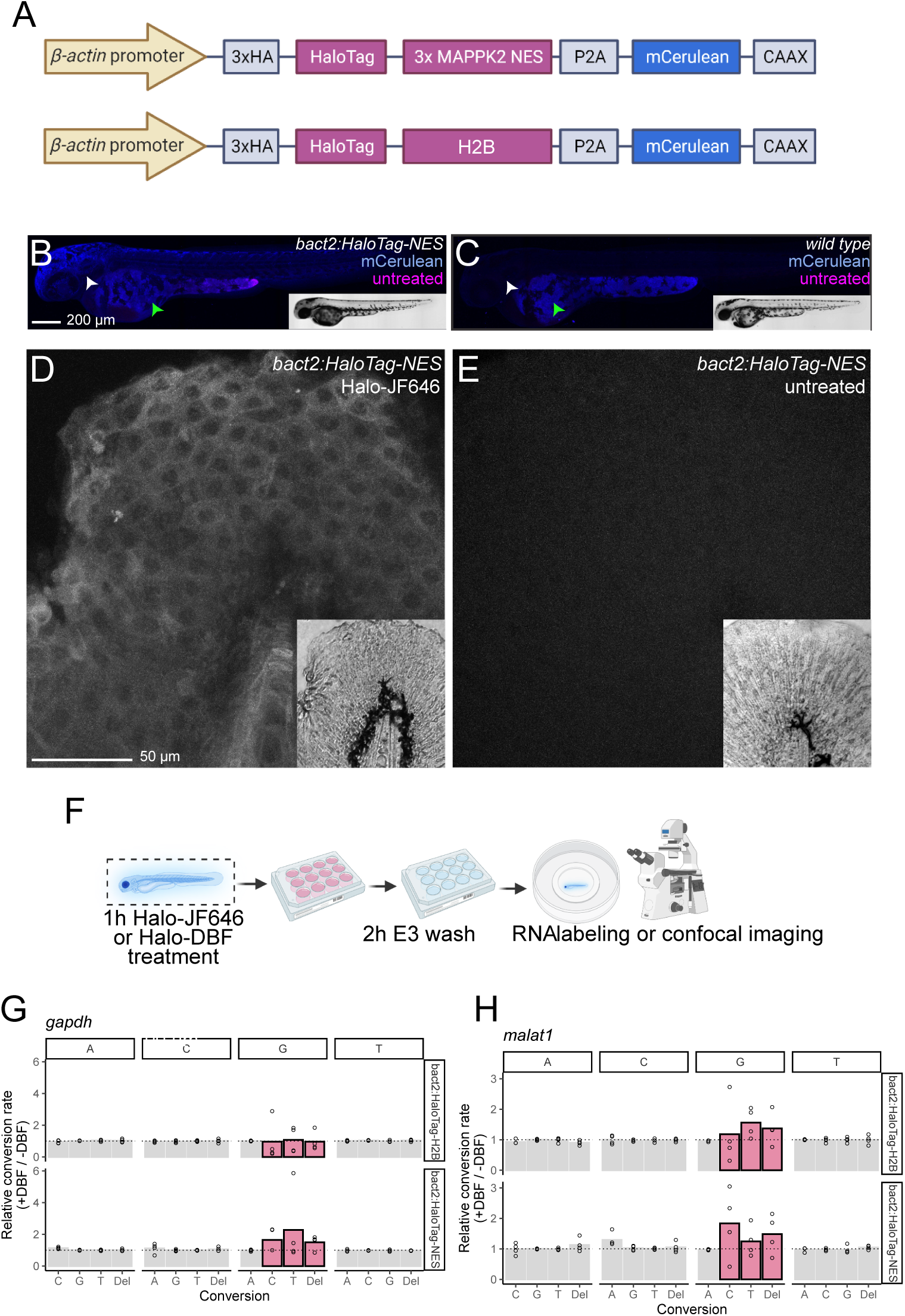
(A) Top: Schematic of the transgenic *bact2:HaloTag-NES-2A-mCerulean-CAAX* zebrafish construct, afterwards referred to as *bact2:HaloTag-NES*. Bottom: Schematic of the transgenic *bact2:HaloTag-H2B-2A-mCerulean-CAAX* zebrafish construct, afterwards referred to as *bact2:HaloTag-H2B*. (B, C) Representative fluorescence imaging of 2 dpf zebrafish embryos either transgenic (B) or negative (C) for the *bact2:HaloTag-NES* transgene. The embryos shown have not been treated with fluorescent Halo ligands. Expression of the HaloTag-NES fusion protein is visualized by mCerulean fluorescence (blue) only in the somatic tissue (white arrow), with background fluorescence seen in the yolk (green arrow, compared to transgene-negative embryo). (D, E) Representative 40x/1 zoom of 2 dpf zebrafish embryo tails of individuals carrying the *bact2:HaloTag-NES* transgene. Zebrafish were exposed to the Halo ligand JF646 (D) or left untreated (E). JF646 staining is seen only in zebrafish exposed to Halo-JF646 (grayscale). (F) Strategy for zebrafish imaging and OINC-seq experiments. (G) Differences in G conversion rates in a *gapdh* amplicon between Halo-DBF treated and untreated samples. Zebrafish genotypes are indicated on the right. (H) As in G, but for an amplicon of *malat1*.

## Notes

### Competing Interest Statement

The authors have declared no competing interest.

## REFERENCES

1 Long, R.M., Singer, R.H., Meng, X., Gonzalez, I., Nasmyth, K. and Jansen, R.P. (1997) Mating type switching in yeast controlled by asymmetric localization of ASH1 mRNA. Science, 277, 383–387.

2 Lécuyer, E., Yoshida, H., Parthasarathy, N., Alm, C., Babak, T., Cerovina, T., Hughes, T.R., Tomancak, P. and Krause, H.M. (2007) Global analysis of mRNA localization reveals a prominent role in organizing cellular architecture and function. Cell, 131, 174–187.

3 Moor, A.E., Golan, M., Massasa, E.E., Lemze, D., Weizman, T., Shenhav, R., Baydatch, S., Mizrahi, O., Winkler, R., Golani, O., et al. (2017) Global mRNA polarization regulates translation efficiency in the intestinal epithelium. Science.

4 Cajigas, I.J., Tushev, G., Will, T.J., tom Dieck, S., Fuerst, N. and Schuman, E.M. (2012) The local transcriptome in the synaptic neuropil revealed by deep sequencing and high-resolution imaging. Neuron, 74, 453–466.

5 Engel, K.L., Arora, A., Goering, R., Lo, H.-Y.G. and Taliaferro, J.M. (2020) Mechanisms and consequences of subcellular RNA localization across diverse cell types. Traffic, 21, 404–418.

6 Ephrussi, A. and Lehmann, R. (1992) Induction of germ cell formation by oskar. Nature, 358, 387–392.

7 Wang, E.T., Taliaferro, J.M., Lee, J.-A., Sudhakaran, I.P., Rossoll, W., Gross, C., Moss, K.R. and Bassell, G.J. (2016) Dysregulation of mRNA Localization and Translation in Genetic Disease. Journal of Neuroscience, 36, 11418–11426.

8 Chen, K.H., Boettiger, A.N., Moffitt, J.R., Wang, S. and Zhuang, X. (2015) RNA imaging. Spatially resolved, highly multiplexed RNA profiling in single cells. Science, 348, aaa6090.

9 Shah, S., Lubeck, E., Zhou, W. and Cai, L. (2016) In Situ Transcription Profiling of Single Cells Reveals Spatial Organization of Cells in the Mouse Hippocampus. Neuron, 92, 342–357.

10. Raj, A. and van Oudenaarden, A. (2009) Single-molecule approaches to stochastic gene expression. Annu. Rev. Biophys., 38, 255–270.

11 Arora, A., Goering, R., Lo, H.-Y. and Taliaferro, J.M. (2021) Mechanical fractionation of cultured neuronal cells into cell body and neurite fractions. Bio Protoc., 11.

12 Taliaferro, J.M., Vidaki, M., Oliveira, R., Olson, S., Zhan, L., Saxena, T., Wang, E.T., Graveley, B.R., Gertler, F.B., Swanson, M.S., et al. (2016) Distal Alternative Last Exons Localize mRNAs to Neural Projections. Mol. Cell, 61, 821–833.

13 Goering, R., Hudish, L.I., Guzman, B.B., Raj, N., Bassell, G.J., Russ, H.A., Dominguez, D. and Taliaferro, J.M. (2020) FMRP promotes RNA localization to neuronal projections through interactions between its RGG domain and G-quadruplex RNA sequences. Elife, 9.

14 Zappulo, A., Van Den Bruck, D., Ciolli Mattioli, C., Franke, V., Imami, K., McShane, E., Moreno-Estelles, M., Calviello, L., Filipchyk, A., Peguero-Sanchez, E., et al. (2017) RNA localization is a key determinant of neurite-enriched proteome. Nat. Commun., 8.

15 Arora, A., Castro-Gutierrez, R., Moffatt, C., Eletto, D., Becker, R., Brown, M., Moor, A.E., Russ, H.A. and Taliaferro, J.M. (2022) High-throughput identification of RNA localization elements in neuronal cells. Nucleic Acids Res., 10.1093/nar/gkac763.

16 Mikl, M., Eletto, D., Nijim, M., Lee, M., Lafzi, A., Mhamedi, F., David, O., Sain, S.B., Handler, K. and Moor, A.E. (2022) A massively parallel reporter assay reveals focused and broadly encoded RNA localization signals in neurons. Nucleic Acids Res., 50, 10643–10664.

17 Mendonsa, S., von Kügelgen, N., Dantsuji, S., Ron, M., Breimann, L., Baranovskii, A., Lödige, I., Kirchner, M., Fischer, M., Zerna, N., et al. (2023) Massively parallel identification of mRNA localization elements in primary cortical neurons. Nat. Neurosci., 26, 394–405.

18 Goering, R., Arora, A., Pockalny, M.C. and Taliaferro, J.M. (2023) RNA localization mechanisms transcend cell morphology. Elife, 12, e80040.

19 Benoit Bouvrette, L.P., Cody, N.A.L., Bergalet, J., Lefebvre, F.A., Diot, C., Wang, X., Blanchette, M. and Lécuyer, E. (2018) CeFra-seq reveals broad asymmetric mRNA and noncoding RNA distribution profiles in Drosophila and human cells. RNA, 24, 98–113.

20 Adekunle, D.A. and Wang, E.T. (2020) Transcriptome-wide organization of subcellular microenvironments revealed by ATLAS-Seq. Nucleic Acids Res., 48, 5859–5872.

21 Padrón, A., Iwasaki, S. and Ingolia, N.T. (2019) Proximity RNA Labeling by APEX-Seq Reveals the Organization of Translation Initiation Complexes and Repressive RNA Granules. Mol. Cell, 75, 875–887.e5.

22 Wang, P., Tang, W., Li, Z., Zou, Z., Zhou, Y., Li, R., Xiong, T., Wang, J. and Zou, P. (2019) Mapping spatial transcriptome with light-activated proximity-dependent RNA labeling. Nat. Chem. Biol., 10.1038/s41589-019-0368-5.

23 Engel, K.L., Lo, H.-Y.G., Goering, R., Li, Y., Spitale, R.C. and Taliaferro, J.M. (2021) Analysis of subcellular transcriptomes by RNA proximity labeling with Halo-seq. Nucleic Acids Res., 10.1093/nar/gkab1185.

24 Fazal, F.M., Han, S., Parker, K.R., Kaewsapsak, P., Xu, J., Boettiger, A.N., Chang, H.Y. and Ting, A.Y. (2019) Atlas of Subcellular RNA Localization Revealed by APEX-Seq. Cell, 178, 473–490.e26.

25 Lo, H.-Y.G., Engel, K.L., Goering, R., Li, Y., Spitale, R.C. and Taliaferro, J.M. (2022) Halo-seq: An RNA Proximity Labeling Method for the Isolation and Analysis of Subcellular RNA Populations. Curr Protoc, 2, e424.

26 Los, G.V., Encell, L.P., McDougall, M.G., Hartzell, D.D., Karassina, N., Zimprich, C., Wood, M.G., Learish, R., Ohana, R.F., Urh, M., et al. (2008) HaloTag: a novel protein labeling technology for cell imaging and protein analysis. ACS Chem. Biol., 3, 373–382.

27 Hein, C.D., Liu, X.M. and Wang, D. (2008) Click chemistry, a powerful tool for pharmaceutical sciences. Pharm. Res., 25, 2216–2230.

28 Steenken, S. and Jovanovic, S.V. (1997) How Easily Oxidizable Is DNA? One-Electron Reduction Potentials of Adenosine and Guanosine Radicals in Aqueous Solution. J. Am. Chem. Soc., 119, 617–618.

29 Alenko, A., Fleming, A.M. and Burrows, C.J. (2017) Reverse Transcription Past Products of Guanine Oxidation in RNA Leads to Insertion of A and C opposite 8-Oxo-7, 8-dihydroguanine and A and G opposite 5-Guanidinohydantoin and Spiroiminodihydantoin Diastereomers. Biochemistry, 56, 5053–5064.

30 Patro, R., Duggal, G., Love, M.I., Irizarry, R.A. and Kingsford, C. (2017) Salmon provides fast and bias-aware quantification of transcript expression. Nat. Methods, 14, 417–419.

31 Pfeiffer, F., Gröber, C., Blank, M., Händler, K., Beyer, M., Schultze, J.L. and Mayer, G. (2018) Systematic evaluation of error rates and causes in short samples in next-generation sequencing. Sci. Rep., 8, 10950.

32 Gonzalez-Rivera, J.C., Sherman, M.W., Wang, D.S., Chuvalo-Abraham, J.C.L., Hildebrandt Ruiz, L. and Contreras, L.M. (2020) RNA oxidation in chromatin modification and DNA-damage response following exposure to formaldehyde. Sci. Rep., 10, 16545.

33 West, J.A., Davis, C.P., Sunwoo, H., Simon, M.D., Sadreyev, R.I., Wang, P.I., Tolstorukov, M.Y. and Kingston, R.E. (2014) The Long Noncoding RNAs NEAT1 and MALAT1 Bind Active Chromatin Sites. Mol. Cell, 55, 791–802.

34 Moll, P., Ante, M., Seitz, A. and Reda, T. (2014) QuantSeq 3′ mRNA sequencing for RNA quantification. Nat. Methods, 11, i–iii.

35 Kaewsapsak, P., Shechner, D.M., Mallard, W., Rinn, J.L. and Ting, A.Y. (2017) Live-cell mapping of organelle-associated RNAs via proximity biotinylation combined with protein-RNA crosslinking. Elife, 6, e29224.

36 Seth, R.B., Sun, L., Ea, C.-K. and Chen, Z.J. (2005) Identification and characterization of MAVS, a mitochondrial antiviral signaling protein that activates NF-kappaB and IRF 3. Cell, 122, 669–682.

37 Schmidt, O., Pfanner, N. and Meisinger, C. (2010) Mitochondrial protein import: from proteomics to functional mechanisms. Nat. Rev. Mol. Cell Biol., 11, 655–667.

38 Li, Y., Aggarwal, M.B., Ke, K., Nguyen, K. and Spitale, R.C. (2018) Improved Analysis of RNA Localization by Spatially Restricted Oxidation of RNA-Protein Complexes. Biochemistry, 57, 1577–1581.

39 Rosenberg, M., Levy, V., Maier, V.K., Kesner, B., Blum, R. and Lee, J.T. (2021) Denaturing cross-linking immunoprecipitation to identify footprints for RNA-binding proteins. STAR Protoc, 2, 100819.

40 Li, Z., Chen, X., Liu, Z., Ye, W., Li, L., Qian, L., Ding, H., Li, P. and Aung, L.H.H. (2020) Recent Advances: Molecular Mechanism of RNA Oxidation and Its Role in Various Diseases. Front Mol Biosci, 7, 184.

41 Simms, C.L., Hudson, B.H., Mosior, J.W., Rangwala, A.S. and Zaher, H.S. (2014) An active role for the ribosome in determining the fate of oxidized mRNA. Cell Rep., 9, 1256–1264.

42 Phillips, C.N., Schowe, S., Langeberg, C.J., Siddique, N., Chapman, E.G. and Resendiz, M.J.E. (2021) Processing of RNA Containing 8-Oxo-7, 8-Dihydroguanosine (8-oxoG) by the Exoribonuclease Xrn-1. Front Mol Biosci, 8, 780315.

43 Khandelia, P., Yap, K. and Makeyev, E.V. (2011) Streamlined platform for short hairpin RNA interference and transgenesis in cultured mammalian cells. Proc. Natl. Acad. Sci. U. S. A., 108, 12799–12804.

44 Schindelin, J., Arganda-Carreras, I., Frise, E., Kaynig, V., Longair, M., Pietzsch, T., Preibisch, S., Rueden, C., Saalfeld, S., Schmid, B., et al. (2012) Fiji: an open-source platform for biological-image analysis. Nat. Methods, 9, 676–682.

45 Schowe, S.W., Langeberg, C.J., Chapman, E.G., Brown, K. and Resendiz, M.J.E. (2022) Identification of RNA Fragments Resulting from Enzymatic Degradation using MALDI-TOF Mass Spectrometry. J. Vis. Exp., 10.3791/63720.

46 Martin, M. (2011) Cutadapt removes adapter sequences from high-throughput sequencing reads. EMBnet journal, 17, 10–12.

47 Dobin, A., Davis, C.A., Schlesinger, F., Drenkow, J., Zaleski, C., Jha, S., Batut, P., Chaisson, M. and Gingeras, T.R. (2013) STAR: ultrafast universal RNA-seq aligner. Bioinformatics, 29, 15–21.

48 Koboldt, D.C., Chen, K., Wylie, T., Larson, D.E., McLellan, M.D., Mardis, E.R., Weinstock, G.M., Wilson, R.K. and Ding, L. (2009) VarScan: variant detection in massively parallel sequencing of individual and pooled samples. Bioinformatics, 25, 2283–2285.

49 Benjamini, Y. and Hochberg, Y. (1995) Controlling the False Discovery Rate: A Practical and Powerful Approach to Multiple Testing. J. R. Stat. Soc. Series B Stat. Methodol., 57, 289–300.

50 Kimmel, C.B., Ballard, W.W., Kimmel, S.R., Ullmann, B. and Schilling, T.F. (1995) Stages of embryonic development of the zebrafish. Dev. Dyn., 203, 253–310.

51 Higashijima, S., Okamoto, H., Ueno, N., Hotta, Y. and Eguchi, G. (1997) High-frequency generation of transgenic zebrafish which reliably express GFP in whole muscles or the whole body by using promoters of zebrafish origin. Dev. Biol., 192, 289–299.

52 Kwan, K.M., Fujimoto, E., Grabher, C., Mangum, B.D., Hardy, M.E., Campbell, D.S., Parant, J.M., Yost, H.J., Kanki, J.P. and Chien, C.-B. (2007) The Tol2kit: a multisite gateway-based construction kit for Tol2 transposon transgenesis constructs. Dev. Dyn., 236, 3088–3099.

53 Mosimann, C. (2022) Multisite Gateway Calculations: Excel spreadsheet. protocolsIO, 10.17504/protocols.io.b4xdqxi6.

54 Felker, A. and Mosimann, C. (2016) Contemporary zebrafish transgenesis with Tol2 and application for Cre/lox recombination experiments. Methods Cell Biol., 135, 219–244.

